# Structural basis of Yta7 ATPase-mediated nucleosome disassembly

**DOI:** 10.1101/2022.05.13.491901

**Authors:** Feng Wang, Xiang Feng, Qing He, Hua Li, Huilin Li

**Affiliations:** Department of Structural Biology, Van Andel Institute, Grand Rapids, Michigan, USA

## Abstract

Yta7 is a novel chromatin remodeler harboring a histone-interacting bromodomain (BRD) and two AAA+ modules. It is not well understood how Yta7 recognizes and unfolds histone H3 to promote nucleosome disassembly for DNA replication. By cryo-EM analysis, we here show that Yta7 assembles a three-tiered hexamer ring with a top spiral, a middle AAA1-tier, and a bottom AAA2-tier. Unexpectedly, the Yta7 BRD stabilizes a four-stranded β-helix termed BRD- interacting motif (BIM) of the largely disordered N-terminal region, and they together assemble the spiral structure on top of the hexamer to engage the nucleosome. We found that the Yta7 BRD lacks key residues involved in acetylated peptide recognition, and as such, it is a noncanonical BRD that does not distinguish the H3 acetylation state, consistent with its role in general DNA replication. Upon nucleosome binding, the BRD/BIM spiral transitions into a flat ring to allow threading of the histone H3 tail into the AAA+ chamber. The H3 peptide is stabilized by the AAA1 pore loops 1 and 2 that spiral around the peptide. Therefore, Yta7 unfolds the nucleosome by pulling on the H3 peptide in a rotary staircase mechanism. Our study sheds light on the nucleosome recognition and unfolding mechanism of Yta7.

## INTRODUCTION

Nucleosomes are a formidable barrier to transcription and other DNA-dependent processes in eukaryotes ^1,2^. A set of chromatin-related factors have evolved to disassemble nucleosomes to facilitate DNA replication, transcription, DNA damage detection, and repair ^3-6^. The *S. cerevisiae* Yta7 (Yeast Tat [trans-activator of transcription]-binding Analog 7) is a chromatin-associated AAA+ (ATPases associated with various cellular activities) ATPase that maintains the balance of nucleosome density of the chromatin ^7-11^. Yta7 is a type II ATPase containing two ATPase domains (AAA1 and AAA2) ^12^ and is conserved in eukaryotes ^13^. The human homolog ATAD2 increases chromatin dynamics and gene transcription, and is frequently overexpressed in various cancers ^14^. The *S. pombe* homologue Abo1 has also been shown to act on nucleosomes ^15^. Yta7, ATAD2, and Abo1 are the only AAA+ ATPase chaperones that acts specifically on histones to regulate gene transcription and DNA replication ^9,13^.

Yta7 was recently found to be a nucleosome segregase that promotes nucleosome disassembly and chromatin replication in an S phase cyclin-dependent kinase (S-CDK) manner ^16^ (**Fig. 1a**). Yta7 was also shown to disassemble the Cse4/H4 tetramer and hand over the unfolded product to Scm3 for deposition to the centromere ^17^. Yta7 contains two tandem AAA+ domains (AAA1 and AAA2) that each contains an α/β subdomain and a helical subdomain and is expected to assemble into a 910-kDa hexamer ^11,15,17^ (**Fig. 1b**). The N-terminal region (1–416 aa) is largely disordered with an acidic N-terminal region (ANR; 118-185 aa) and 13 phosphorylation sites ^13^, but likely contribute to the H3 recognition although the underlying mechanism is unclear ^18^. The C-terminal region contains an extended bromodomain (BRD). BRD is a conserved protein-interacting module that recognizes histones and regulates ATP-dependent chromatin remodeling ^19-21^. A typical BRD fold is 110 residues long and composed of a four-helix bundle that binds to acetylated peptides. Interestingly, the extended Yta7 BRD binds the H3 N-terminal peptide (N-tail) regardless of acetylation status ^18^. Although the structure of several canonical BRDs have been determined ^22^, and in fact BRDs are being targeted for cancer drug development ^19,23-26^, how the Yta7 BRD recognizes histone H3 is unknown ^18,27^.

**Figure 1.**
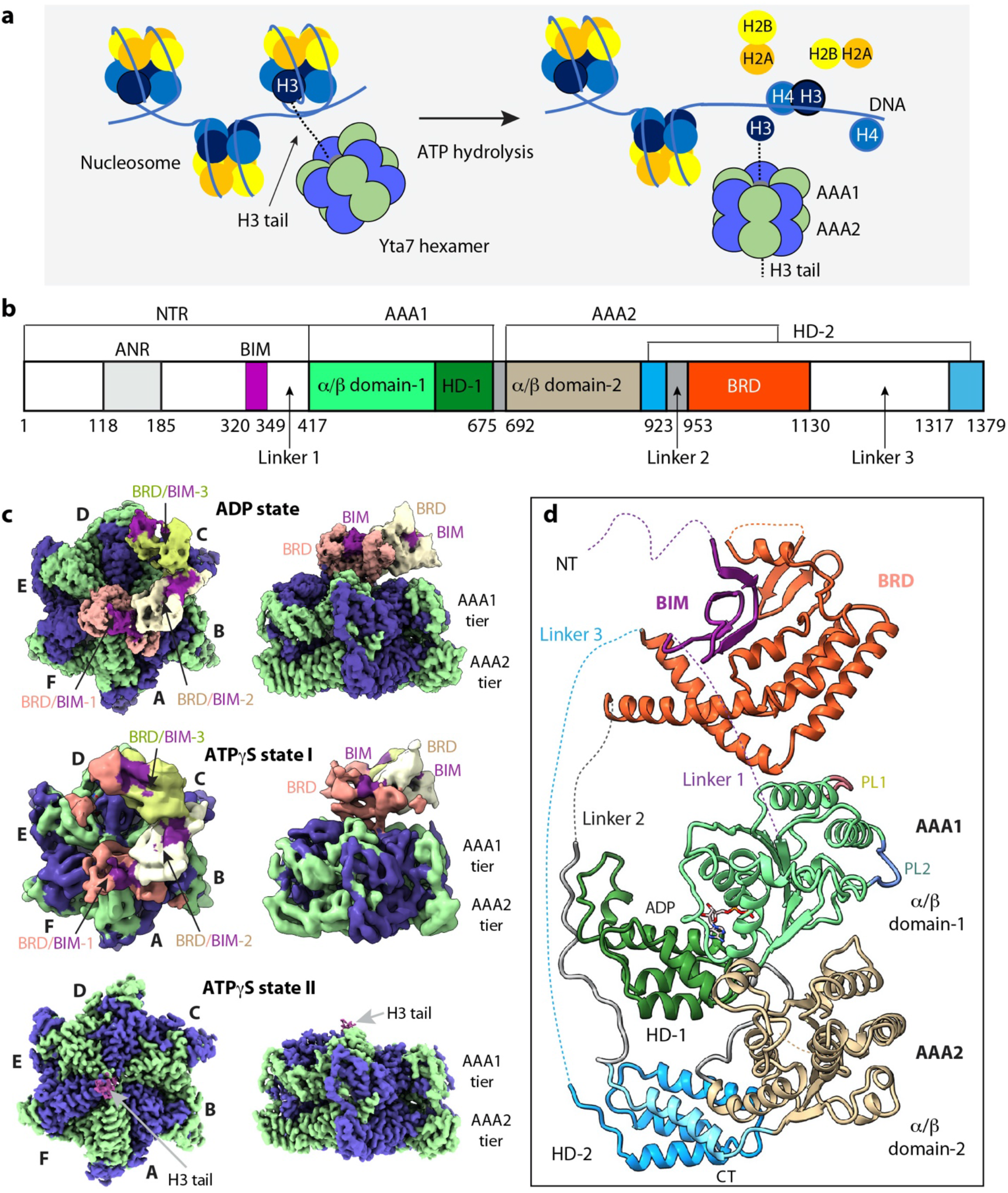
Structures of the *S. cerevisiae* Yta7 ATPase hexamer. **a**, A sketch showing that Yta7 unfolds histone H3 to remove nucleosomes from chromatin. **b**, Domain architecture of the yeast Yta7. Unresolved regions in the EM map are in white. **c**, Top and side views of the EM map of the Yta7 hexamer bound to ADP or ATPγS in states I and II. Alternating subunits are in blue and green. The three resolved BRDs are in salmon, cream, and olive, respectively, and the associated BIMs are in magenta. The H3 tail is in plum. **d**, Structure of an Yta7 protomer with individual domains colored as in (b). NT, N-terminus, CT, C-terminus.

The first structural insight was obtained by a recent structural study of the *S. pombe* homologue Abo1, revealing stacked AAA1 and AAA2 rings, with the rings adopting either an open spiral or a closed configuration depending on the nucleotides ^15^. However, the study used an N-terminal region truncated protein, did not model the BRD, and captured a translocating peptide of unknown source. Therefore, the molecular mechanism underlying histone recognition and unfolding by Yta7 is still unclear. We have performed cryo-EM analysis on the purified full-length Yta7 and obtained structures of Yta7 with ATPγS at 3.0-Å resolution and with ADP at 3.1-Å resolution, providing the first insight into the BRD arrangements on top of the AAA+ rings. We found that the Yta7 BRD stabilizes a BRD-interacting motif (BIM) of the N-terminal region, and the BIM promotes BRDs oligomerization and contributes to nucleosome recognition. The structure explains why the BRD does not distinguish the H3 peptide acetylation and suggests a hand-over-hand mechanism for H3 unfolding. We further determined an 18-Å EM map of the Yta7–nucleosome complex, which suggests that the nucleosome is held on the top of the BRD ring by the Yta7 N-terminal region. Our study provides insights into nucleosome recognition and the unfolding mechanism of Yta7 and contributes to understanding of the nucleosome disassembly.

## RESULTS

### 1. Yta7 structure

We first purified the full-length Yta7 and determined the cryo-EM structure in the presence of 2 mM ADP at an average resolution of 3.1 Å (**Fig. 1c-d, Supplementary Figures 1-2**). The ADP-bound structure was resolved with three ordered BRD domains forming a spiral above the AAA1 tier (**Supplementary Figure 3**). To understand how Yta7 interacts with the H3 tail, we next performed cryo-EM on the Yta7–H3 tail (1-24 aa) complex in the presence of 2 mM ATPγS. We derived two 3D maps; one resembled the ADP-bound structure with three ordered BRD domains in a right-handed spiral form (state I) at an estimated resolution of 5.6 Å and the other with a bound H3 tail at 3.0 Å resolution (state II) (**Fig. 1c, Supplementary Figures 4-6**). The three long linkers (Linkers 1-3) are expected to be flexible and indeed their densities were largely missing except for the first half of Linker 2 (**Fig. 1d**). As expected, Yta7 assembled a hexamer similar to the pombe homolog Abo1 ^15^. In the ADP-bound Yta7 hexamer, we were able to resolve three out of the six BRDs at a lower resolution of 3.9 Å (**Supplementary Figure 1a**). The structure is three-tiered with a spiral BRD-tier on the top followed by the AAA1 and AAA2 tiers, has a dimension of 150 × 150 × 135 Å, and adopts an asymmetrical overall configuration (**Fig. 1c-d, Supplementary Figure 3a**). The six AAA2 domains form a flat and symmetric ring at the bottom tier. But the six AAA1 domains form a shallow spiral ring in the middle tier, enabled by the six well-ordered but differently configured linkers between the AAA1 AAA2 domains (**Supplementary Figure 3b**). Unexpectedly, we identified a 25-residue motif from the N-terminal region (Gly320-Asn349) in the BRD region based on the density feature and the AlphaFold structural prediction ^28^, and have referred to it as a BRD-interacting motif (BIM) (**Fig. 1c-d)**. The BIM is not conserved and appears to be unique to Yta7 (**Supplementary Figure 7**).

**Figure 2.**
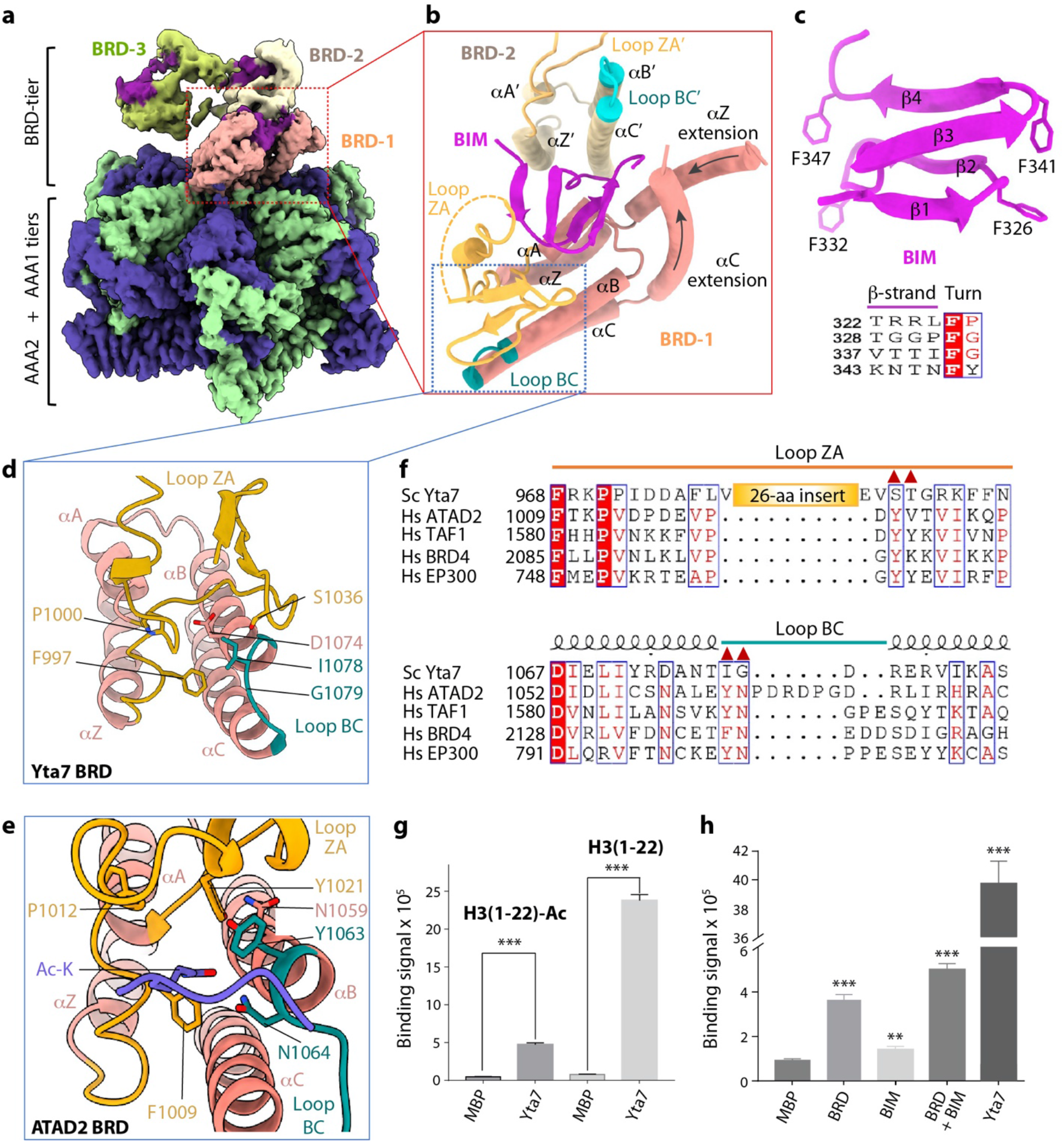
Structure of the noncanonical Yta7 bromodomain and the novel BRD- interacting motif (BIM). **a**, Surface rendered composite EM map of the ADP-bound Yta7 hexamer colored in the same scheme as in Fig. 1d. The EM densities of the 2nd (in cream) and 3rd BRD (in olive) and their associated BIMs (in magenta) were derived separately and surface- rendered at a lower threshold. **b**, Interface structure between BRD/BIM-1 and BRD/BIM-2. The conserved α-helices (αA-αC, αZ) and their connecting loops (ZA in yellow and BC in teal) of the BRD fold are labeled. **c**, Structure of the BIM motif in cartoon. Lower panel shows the conserved Phe at each turn of the β-strands. **d-e**, Comparison of the substrate-binding pocket of BRD in Yta7 (d) and in ATAD2 (e; PDB 4QUT). Key residues involved in H3 N-tail binding are labeled. **f**, Sequence alignment of the ZA and BC loops of five BRDs. The red triangles mark the conserved residues that bind to the acetylated lysine in ATAD2 and PCAF; these residues are not conserved in Yta7. Sc, *Saccharomyces cerevisiae;* Hs, *Homo sapiens*. **g**, Interaction between Yta7 and unmodified and acetylated H3 N-tails (1-22 aa). All values represent means ± SD obtained from five independent experiments performed in triplicate. **h**, Interaction of the H3 N-tail (1-22 aa, biotinylated) either with BRD alone (with N-terminal His-tag and MBP fusion) or with NTR alone (with N-terminal His-tag and MBP fusion), or with the BrD–NTR complex (both are fused with MBP). The 6xHis-MBP served as a negative control (n = 3; error bars = SD), *** significantly different from control (P ≤ 0.001, one-tailed t-test), ** (P ≤ 0.01, one- tailed t-test).

Corresponding to the three-tiered architecture of the hexamer, a Yta7 subunit can be divided into three main regions: the AAA1 and AAA2 at the middle and bottom regions, and the top region comprised of the N-terminal BIM and the C-terminal BRD (**Fig. 1d**). This organization places the unresolved N-terminal 320 residues on the top of the Yta7 hexamer where the nucleosome engages, suggesting that the disordered N-terminal region with its associated ANR may play a role in nucleosome recognition. AAA1 is a canonical AAA+ domain with a nucleotide binding pocket between the α/β subdomain and the helical subdomain and containing the ATPase activity. AAA2 adopts an inactive α/β subdomain without a nucleotide binding site and its helical subdomain is a composite of one α-helix following the α/β-subdomain and 4 α-helices at the C-terminus following the BRD. Therefore, the BRD appears to be “inserted” into the AAA2 sequence but is projected all the way up on the top of the structure by the two long loops – the 37-residue linker 2 and the 187-residue linker 3 (**Fig. 1d**). This architecture contrasts with the general disaggregase ClpB/Hsp104 in which an essential middle domain is inserted in the AAA1 tier and is located to the side of the hexamer ^12,29,30^.

### 2. Both BRD and BIM contribute to histone H3 binding

The most striking feature in the ADP-bound Yta7 is the BRD/BIM spiral rising above the AAA1 tier. The proximal BRD/BIM-1 is from subunit B but interacts with the top AAA1 surface of subunits A, F, and E and blocks the entry into the AAA+ chamber (**Figs. 1c, 2a, Supplementary Figure 3a**). The Yta7 BRD spans from Pro954 to Leu1136 and contains the conserved core of the four-helix bundle (αZ, αA-αC) (**Fig. 2b**). However, the core is expanded by one α-helix preceding the αZ (αZ extension) and another α-helix following αC (αC extension). The expansion is unique to Yta7 and ATAD2 and is absent in a canonical BRD. The BRD- interacting motif (BIM) is cradled in the U-shaped BRD burying a surface of 890 Å^2^, over 30% of the total surface, and the BIM has a right-handed 2-turn β-helix fold, a fold frequently involved in protein-protein interaction ^31^ (**Fig. 2b**). Interestingly, at each turn of the four β-strands is located a hydrophobic phenylalanine residue (**Fig. 2c**). The Yta7 BIM mediates and cements the inter-subunit BRD-BRD interaction. Therefore, Yta7 uses its N-terminal BIM to position and stabilize the BRD spiral on top of the structure. In the absence of the BIM, the BRD spiral would not be able to reside on top of the AAA1 tier, because the BRD is inserted in the AAA2 that is at the bottom of the hexamer and there is little direct BRD-BRD interaction.

The ZA loop of Yta7 BRD has a 26-residue insertion compared to canonical BRDs (**Fig. 2d-f**). The ZA loop is stabilized by BIM. In the canonical BRDs, the ZA and BC loops form a pocket that specifically recognizes acetylated histone peptides ^20,22^. But the longer Yta7 ZA loop makes the substrate binding pocket more open (**Fig. 2d**). Furthermore, the conserved Tyr in the ZA loop of human ATAD2 and other BRD-containing proteins is replaced by Ser in Yta7, and the conserved Tyr/Phe-Asn dipeptide motif in the BC loop is replaced by Ile-Gly in Yta7 (**Fig. 2e-f**). These substitutions likely account for the observation that Yta7 binds to histones in a posttranslational modification-independent manner ^18^. We next examined Yta7’s binding affinity with the H3 tail (1-22 aa) either in the non-acetylated or in the acetylated form by AlphaScreen (Amplified Luminescent Proximity Homogeneous Assay). In the acetylated form of H3 tail, all four lysine residues (Lys4, Lys9, Lys14, and Lys18) were acetylated. We found that the binding signal of the nonacetylated H3 N-tail was 5 times stronger than that of acetylated H3 tail (**Fig. 2g**), confirming our structural based observation.

Because BIM is near the BRD substrate binding pocket and stabilizes the BRD/BIM spiral, we asked if BIM also plays a role in histone binding. We purified 6xHis-MBP-fused BRD (954-1132 aa) and MBP-fused BIM (320-350 aa) with or without 6xHis-tagging, respectively, and used the AlphaScreen assay to examine their binding to the H3 tail (**Supplementary Figure 8**). We found that BIM alone had weak binding signal that was weaker than the BRD alone but stronger than the MBP control (**Fig. 2h**). Furthermore, the binding signal of BIM and BRD together was stronger than the BRD alone. These results suggest that the BIM participates in the BRD recognition of the H3 tail. We note that the binding signal of BRD/BIM is an order of magnitude weaker than that of the Yta7 hexamer, probably because the multiple BRD/BIM domains present in the full-length Yta7 hexamer all contribute to the H3 binding.

### 3. The Yta7 AAA1 pore loops spiral around the histone H3 tail

When the BRD/BIM-tier is in the partially ordered spiral form, as observed in the ADP state and in the ATPγS state I, the proximal BRD/BIM-1 blocks the entry of the AAA+ chamber, and correspondingly, there is no density for the H3 tail in the AAA1/2 chamber (**Fig. 1c**). However, when the BRD/BIM-tier is disordered as found in the ATPγS state II, the substrate entry is apparently open, and we observed a continuous and linear density with side chain features in the central chamber of the Yta7 AAA1 tier. We were able to assign the density to the C-terminal 15 residues (10-24 aa) of the 24-residue H3 tail we used in the experiment (**Figs. 1c, 3a**). The resolved H3 peptide is oriented with its N-terminus pointing downwards (**Fig. 3b**), suggesting that the N-terminus of H3 first threads into the Yta7 central chamber, and that the unresolved N-terminal 9 residues must have reached to the AAA2-tier chamber and are disordered. This is consistent with the fact that AAA2 chamber is larger (24 – 32 Å) than the AAA1 chamber (8 – 14 Å) and lacks the peptide-translocating pore loops present in the AAA1 tier (**Fig. 3a**).

**Figure 3.**
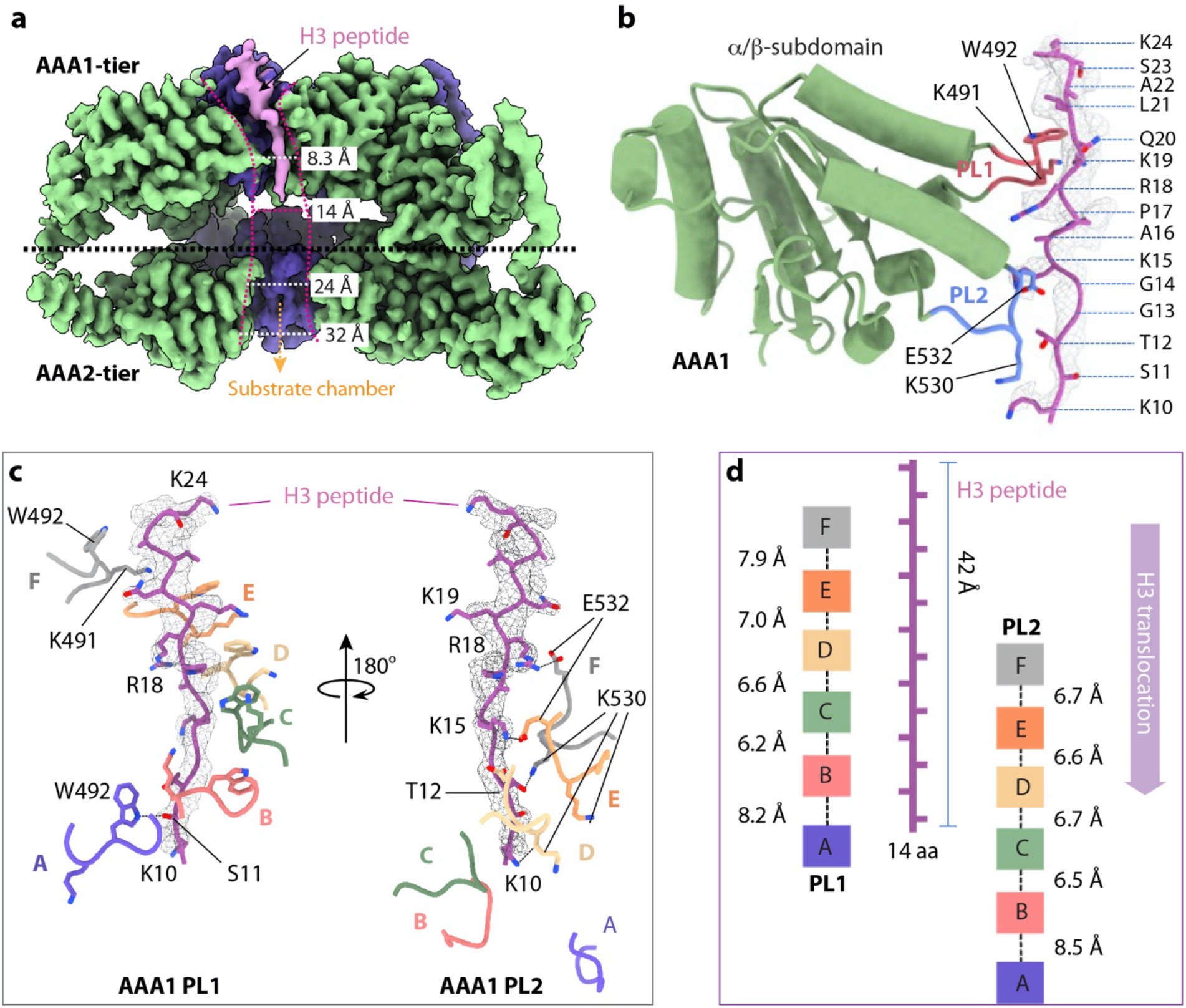
The H3 N-tail coordination in the central chamber of the ATPγS-bound Yta7 hexamer. **a**, A cut-open sideview of the 3D map. The EM density of the trapped H3 N-tail is in salmon. **b**, Structure of the AAA1 α/β sub-domain with its PL1 (yellow) and PL2 (cyan blue) loops coordinating the H3 tail. The H3 tail structure is in sticks superimposed with the EM density in gray meshes. **c**, Close-up views of the interactions between H3 tail and the coordinating PL1(left) and PL2 (right) loops of the AAA1 domain α/β subdomain. The EM density of H3 N-tail superposed and in grey mesh. Trp492 and Lys491 (shown as sticks) from PL1 form a staircase around the H3 peptide. Hydrogen bonds are shown as black dash lines and the involved residues are labeled. The 7-Å axial distance is measured from PL1 Trp492 to PL2 Glu532. **d**, Schematic of the H3-interacting double spiral formed by AAA1 PL1 and PL2 loops, with inter-loop distances along the substrate axis shown based on the position of PL1 Trp492 (left) and PL2 Glu532 (right).

The H3 peptide Lys10-Lys24 is stabilized by pore loops 1 (PL1) and 2 (PL2) of the Yta7 AAA1 α/β subdomain (**Fig. 3b**). Specifically, six PL1 (one from each of the six subunits) surround the peptide like a right-handed staircase with six Trp492 residues contacting the peptide backbone (**Fig. 3c**). However, the lowest Trp492 of subunit A has weaker interaction with the H3 peptide because its side chain points away from the substrate, suggesting that the subunit A PL1 is about to disengage from the substrate. The six PL2 also form a right-handed spiral to surround the substrate peptide, but only four PL2 loops (of subunits C-F) contact the peptide, via hydrogen bonds between subunit F Glu532 and H3 Arg18 and between subunit E Glu532 and H3 Arg15, and between subunit D Lys530 and H3 Lys10. The two PL2 of the lower subunits A and B are away from the H3 peptide. These substrate-interacting residues are conserved in the Abo1 structure. It was found that the Abo1 mutations W345A and E385A – equivalent to the Yta7 W492A and E532A– diminished the histone manipulation function while not affecting the ATPase activity, underscoring the importance of these peptide-binding residues ^15^. The axial distance between the H3-interacting Trp492 in PL1 and Glu532 in PL2 is 7 Å, indicating a peptide translocation step size of two amino acids (**Fig. 3d**). The aromatic pore loops spiraling around the peptide substrate and the two-residue per translocation step are conserved in several well-characterized AAA+ protein unfoldases ^12,29,30^.

### 4. Sequential ATP hydrolysis cycle in the AAA1-tier

The resolution of our cryo-EM maps was sufficient to identify the bound nucleotides (**Fig. 4a-b**). We observed no nucleotide in the AAA2-tier in all Yta7 structures, consistent with the knowledge that the AAA2 is an inactive AAA+ fold. In the ADP-bound Yta7, we identified four ADP molecules in the nucleotide binding pockets of subunits A-D, a partially occupied ADP in subunit E, and no ADP in subunit F (**Fig. 4a**). In the ATPγS-bound Yta7, five ATPγS molecules were identified in subunits B-F. The EM density in the nucleotide binding pocket of the lowest subunit A is consistent with an ADP, suggesting the ATPγS in this site has been hydrolyzed (**Fig. 4a-b**). Interestingly, the entry to the nucleotide binding pocket is largely open in the ADP-bound state but is closed in the ATPγS-bound state (**Fig. 4a**). The entry is controlled by two gating loops: gate loop 1 is formed by Pro403-Asn409, and gate loop 2 is formed by Met549- Arg552 that connects α3 and β4 of an adjacent AAA1. In the ATPγS-bound state, gate loop 1 of subunit D moves towards subunit C and gate loop 2 of subunit C clamps down to cover the entry path. These loops move away from their positions in the ATPγS-bound state to open the nucleotide entry path, perhaps facilitating nucleotide exchange in the ADP state.

**Figure 4.**
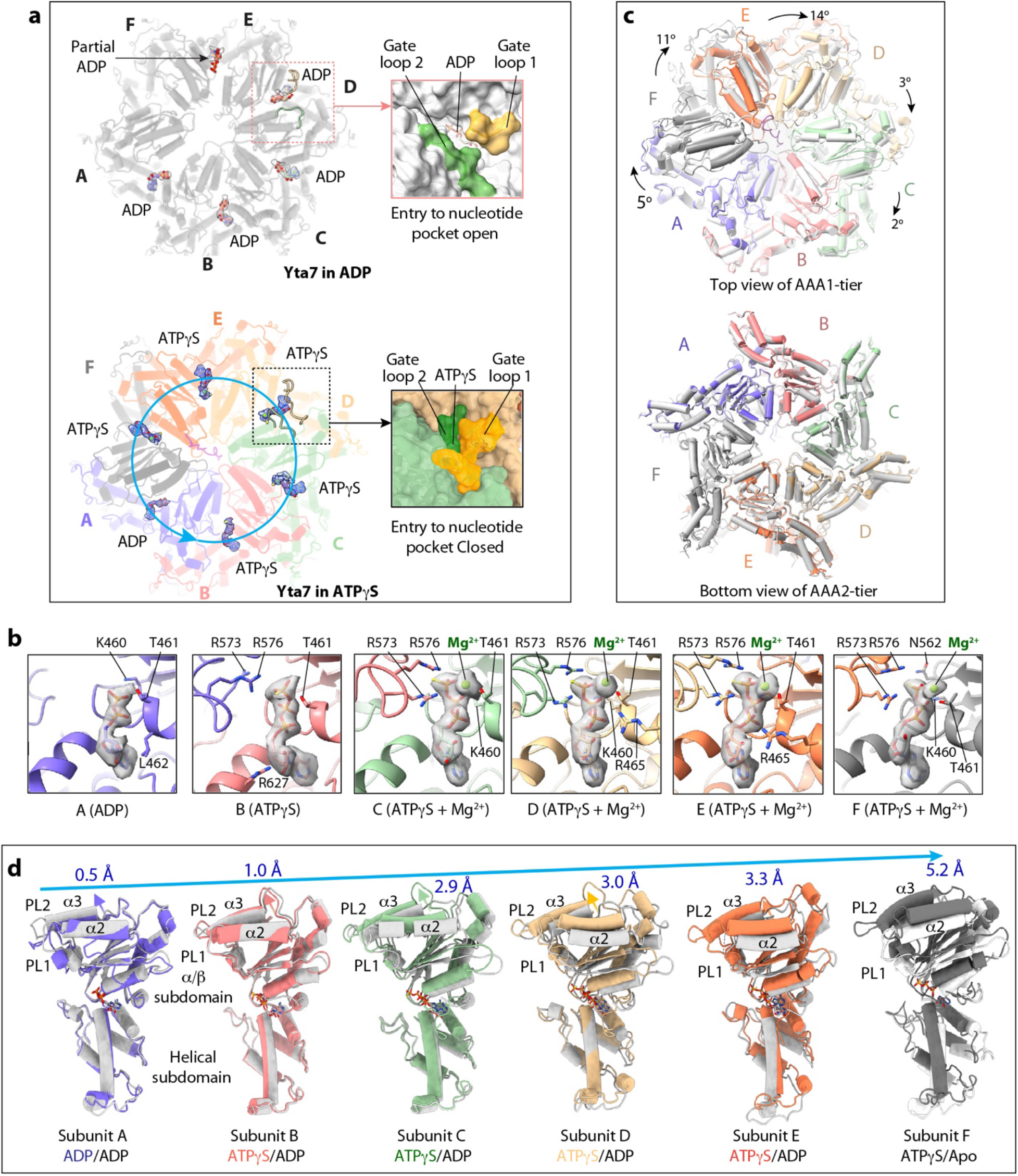
Nucleotide binding in the ADP- and ATPγS-bound Yta7 structures. **a**, Top view of the AAA1-tier of Yta7 in the ADP- and ATPγS-bound state. The bound nucleotides are in sticks superimposed with the EM map rendered in transparent surface views. The arrowed cyan circle indicates the sequence of ATP hydrolysis in the counterclockwise direction. The ADP/ATPγS binding pocket in subunit D of Yta7 in the ADP- and ATPγS-bound states are shown in enlarged views on the right. The nucleotide entry/exit gate is open in the ADP-bound state but is closed by the gate-1 and gate-2 loops in the ATPγS-bound state. **b**, Detailed view of the nucleotide- binding site in each subunit in the ATPγS-bound Yta7. ADP/ATPγS are shown in sticks with their respective EM densities superimposed in transparent gray surface view. The Mg^2+^ ions resolved in subunits C-F are in green spheres. The six EM densities are rendered at the same threshold. The residues that coordinate the nucleotide are in sticks and labeled. **c**, Superposition of the Yta7 AAA+ hexamer in the ADP- (grey) and ATPγS-bound (color) states. Top panel shows the top view of AAA1-tier, and the bottom panel shows the bottom view of AAA2-tier. **d**, Side-by-side comparison of the AAA1/2 region of each subunit in the ADP- and ATPγS-bound states. The PL1 and PL2 loops and the α2, and α3 helices of the AAA1 domain gradually move upward from left to right in the ATPγS-bound state. The arrows indicate the rigid movement of AAA1 from the ADP-bound state to ATPγS-bound state. The movement distance of the α3 helix is labeled on the top of each panel.

In the ATPγS-bound Yta7 structure, the five ATPγS molecules in subunits B-F are stabilized by Lys460 and Thr461 of the hosting subunit and Arg573 and Arg576 of the adjacent subunit (**Fig. 4b**). In subunit A, the ADP is stabilized only by Lys460 and Thr461, and the Arg573 and Arg576 of the adjacent subunit are too far to interact. The Mg^2+^ ions coordinating the ATPγS molecule are resolved in subunits C-F, but not in subunit B, probably due to a lower occupancy of the cation at this location. We expect that ATP hydrolysis occurs counterclockwise in the order of subunit A-B-C-D-E-F-A (**Fig. 4a**). In other word, the next subunit to hydrolyze ATP is B, and when subunit B takes on the pose of subunit A, its pore loops PL1 and PL2 will move down by ∼ 8 Å, thereby pulling the bound H3 peptide downwards by two amino acids (**Fig. 3d**). The six inactive AAA2 domains form a symmetric ring and there is virtually no conformational change between the ADP and ATPγS states (**Fig. 4c**). However, the AAA1 domain undergoes nucleotide-dependent rigid-body movement: these domains rotate individually clockwise by 2°- 14° (**Fig. 4c**) and moves up by 0.5 Å to 5.2 Å from the ADP to ATPγS state (**Fig. 4d**). These structural changes likely occur during the ATP hydrolysis-driven histone unfolding process by the Yta7 hexamer, which is consist with other well-characterized protein unfoldases ^32-37^.

### 5. Nucleosome induces structural rearrangement in Yta7

We next investigated how Yta7 recognizes the nucleosome. We co-expressed the yeast H2A, H2B, H3, and H4 in *E coli*, purified the histone octamer (**Fig. 5a**), and reconstituted by gradient dialysis the yeast nucleosome with 167-bp Widom 601 sequence ^38^. Yta7 bound reconstituted yeast nucleosome very weakly. Therefore, we employed the GraFix technique to stabilize the interaction with 0–0.15% glutaraldehyde ^39^. 2D class averages revealed the largely flexible positioning of the nucleosome on top of the Yta7 (**Fig. 5a, Supplementary Figure 9b-c**).

**Figure 5.**
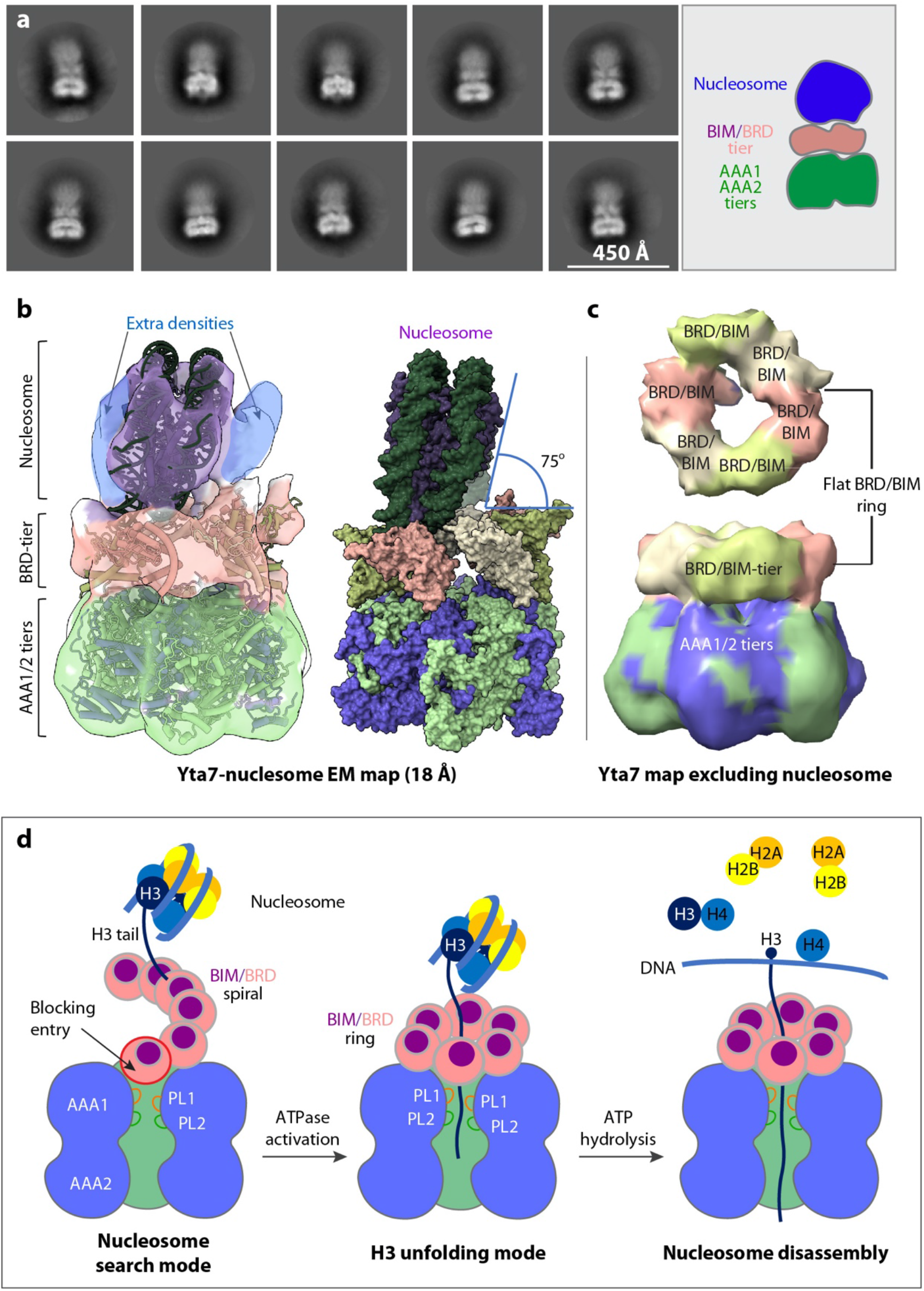
Nucleosome binding and conformational changes of Yta7. **a**, Selected 2D class averages of the Yta7–nucleosome complex. **b**, Cryo-EM map in a transparent surface view, superposed with the structures of Yta7 and nucleosome (PDB 6GEJ) shown in cartoons. Right panel shows the atomic model in surface view. The extra EM densities outside the nucleosome likely belong to the disordered Yta7 N-terminal regions that may participate in nucleosome binding. **c**, Top and side views of the EM map derived by local refinement excluding the nucleosome region. The EM density of the AAA1/2-tiers is omitted in the top view for clarity. The BRD/BIM tier is a flat ring with a central pore leading towards the Yta7 chamber. **d**, Schematic model of the Yta7-mediated nucleosome removal from chromatin. In the initial search mode, the Yta7 BRD/BIM spiral upwards to engage a nucleosome. The entry to the Yta7 central chamber is blocked by the proximal BRD/BIM. Next, the nucleosome engages with Yta7 with its edge. Nucleosome binding coverts the BRD/BIM spiral to a flat ring and unblocks the entry to the Yta7 chamber. When the H3 tail is inserted into the AAA1 chamber, the PL1 and PL2 loops pull on the H3 tail. Yta7 unfolds H3 in a hand-over-hand mechanism driven by ATP hydrolysis. Finally, H3 unfolding leads to nucleosome disassembly.

Because of the varied association of the nucleosome, we used 3D variability analysis to derive a series of 3D reconstructions of the Yta7–nucleosome complex ^40^ (**Supplementary Figure 9d**). Among these, one 3D map at 18-Å resolution had relatively clear nucleosome density (**Fig. 5b, Supplementary Fig. 9**). Rigid-body docking of the nucleosome structure and the Yta7 structure into the EM map reveal that the nucleosome approaches Yta7 edge-on at an angle of ∼75° (**Fig. 5b**). We next performed focused refinement by excluding the flexible nucleosome region, leading to a medium resolution 3D map (14 Å) of the Yta7 hexamer (**Fig. 5c**). The Yta7 structure determined in the presence of nucleosome shows that the BRD/BIM-tier has converted from the spiral shape in the absence of nucleosome into a flat ring, as it is sandwiched between the nucleosome and the AAA1-tier. The nucleosome appears to have reconfigured the BRD/BIM-tier, such that the entry into the AAA1 central chamber is no longer blocked as seen in the absence of nucleosome (**Fig. 5c**). We noticed extra densities on both sides of the nucleosome (**Fig. 5b**). They are probably from the unstructured N-terminal regions of the hexamer, especially the conserved ANR (acidic N-terminal region). This possibility is consistent with the previous suggestion that the long and flexible N-terminal region participates in nucleosome recruitment ^18^.

## DISCUSSION

Yta7 is a recently identified chromatin segregase that disassembles nucleosomes in an S-CDK dependent manner to promote chromosome replication ^16^. We have solved the first nearly full-length structure of Yta7. The previous structural study of the pombe homolog Abo1 used a N-terminal truncated version, and the BRD was not modeled due to low resolution in that region ^15^. We discovered that the Yta7 BRD is not only expanded by two additional α-helices, but also with a much longer ZA loop and a distinct BC loop. These structural features underlie our observation that the Yta7 BRD binds more tightly to unmodified histone H3 tail than to the acetylated version. We identified a novel BRD-interacting motif (BIM) in the N-terminal region with a β-helix fold that stabilizes the BRD spiral on top of the AAA+ hexamer for nucleosome recognition. We further demonstrated that BIM itself contributes to the recognition of the H3 tail.

Importantly, we found that the six BRD/BIM assemble a spiral structure that likely facilitates the search and initial engagement of Yta7 with the nucleosome. Our cryo-EM analysis revealed that the BRD/BIM spiral is converted into a flat ring upon nucleosome binding, and that the flat BRD/BIM ring contains a central pore that allows the H3 tail to thread through. Once it enters Yta7, the H3 peptide is coordinated by the spirally arranged pore loops in the AAA1 chamber. The nucleotide binding pattern and the pore loop arrangement in the peptide-translocating AAA1-tier support a counterclockwise sequential ATP hydrolysis cycle and a processive peptide unfolding mechanism of the Yta7 hexamer. Therefore, the peptide-translocation aspect of the Yta7 is conserved in other protein unfoldases ^34-37^.

Therefore, our study has revealed that the Yta7 hexamer functions in two distinct modes to disassemble the nucleosome (**Fig. 5d**). The first is a nucleosome search mode in which the six BRD/BIM domains assemble a spiral to recruit the histone H3 tail. Although we have demonstrated that both BRD and BIM contribute to the H3 recognition, we do not yet know how this initial recognition occurs, because the H3 peptide visualized in our cryo-EM maps has already entered the AAA1 chamber. Following the nucleosome engagement, Yta7 transits into the second mode by converting the BRD/BIM spiral into a flat ring that allows the H3 tail to thread into the AAA1 chamber. This nucleosome-binding induced transition likely activates the Yta7 ATPase, leading to the processive H3 unfolding and eventually the disassembly of the nucleosome. In summary, our work has shed new light on the Yta7-mediated nucleosome disassembly. Further studies are needed to understand how the disordered N-terminal region contributes to the nucleosome binding, and how the BRD/BIM recognizes the H3 tail before admitting it into the AAA1 chamber for unfolding.

## Acknowledgements

Cryo-EM micrographs were collected at the David Van Andel Advanced Cryo-Electron Microscopy Suite at Van Andel Institute. We thank G. Zhao and X. Meng for facilitating data collection. We thank H. Wen and Z. Xue at Van Andel Institute for providing biotinylated H3 peptide and biotinylated H3 acetylated peptide for the AlphaScreen assay. This work was supported by the US National Institutes of Health grants GM131754 (to Huilin Li) and Van Andel Institute (to Huilin Li).

## Author contributions

F.W. and Huilin Li. designed research; F.W., X.F., Q.H., and Hua Li performed research and analyzed the data; F.W. and Huilin Li wrote the manuscript with input from all authors.

### Competing interests

The authors declare no competing interests.

### Data Availability

The cryo-EM 3D maps of the *S. cerevisiae* Yta7 bound to ADP and Yta7 bound to H3 tail and ATPγS have been deposited in the Electron Microscopy Data Bank with accession codes EMD-26695, EMD-26696, EMD-26697, and EMD-26682. The corresponding atomic models have been deposited in the Protein Data Bank with accession codes 7UQI, 7UQJ, and 7UQK.

## MATERIALS AND METHODS

### DNA constructs generation

*S. cerevisiae YTA7* gene was amplified from yeast DNA genome and cloned into the integrated and galactose-inducible vector pRSII403 with an N-terminal 10xHis tag and 3xFLAG tag. The truncated DNA encoding Yta7 BRD-interacting motif (BIM) (320-350) and BRD (956-1125) were cloned into a pETDuet1 vector (Novagen) in fusion with an N-terminal 6xHis-MBP tandem tag. The Yta7 BIM (320-350) was also cloned into a pET29a expression vector with a MBP fusion but without the 6xHis tag. Genes encoding *S. cerevisiae* histones H2A, H2B, H3 and H4 were cloned into and replaced the same *Xenopus laevis* histone regions in the polycistronic co-expression plasmid pET29a-YS14 (Addgene #66890). All plasmids DNA constructs were verified by sequencing.

### Protein expression and purification

The pRSII403-Yta7 plasmids were linearized and transformed into yeast strain ySK119 (W303 background) ^41^ to integrate in the HIS3 region. The strains and plasmids used in this work are described in **Supplementary Table 1**. Yeast strain overexpressing Yta7 was grown in selective medium before being inoculated into 9 L of the YP-raffinose medium at 30°C. Cells were grown to the OD_600_ of 1.0 and arrested for 3 h with 100 ng/ml of α-factor (GenScript). Then 2% (w/v) galactose was added to induce protein overexpression for 4-5 h at 30°C. After induction, the cells were collected and washed with 200 ml ice-cold 25 mM HEPES-KOH pH 7.6, 1 M sorbitol once and washed twice with lysis buffer (250 mM KCl, 25 mM HEPES-KOH pH 7.6, 10 % glycerol, 0.05% NP-40, 1 mM EDTA, 4 mM MgCl_2_). The cell pellets were resuspended in the same cell volume of the lysis buffer containing two protease inhibitor tablets, and the suspensions were frozen drop-by-drop into liquid nitrogen. The frozen cells were crushed using a freezer mill (SPEX CertiPrep 6850 Freezer Mill) for 12 cycles, 2 min each at 15 cps with 2 min cooling in between. Powder was slowly thawed on ice overnight, then mixed into an equal amount of lysis buffer plus protease inhibitors. Insoluble material was cleared by centrifugation in the Ti-45 rotor at 40,000x rpm for 1 h. The supernatant was incubated in batch with pre-washed anti-FLAG M2 affinity gel (Sigma) for 4 h at 4°C. After flow through, the beads were washed by 50 column volume (CV) lysis buffer, then washed again by another 50 CV lysis buffer. Finally, the proteins were eluted with 5 CV lysis buffer containing 0.3 mg/ml Flag peptides. Yta7 was further purified by size exclusion chromatography using a Superose 6 10/300 GL column (GE Healthcare) in a cold buffer (150 mM KCl, 25 mM HEPES-KOH pH 7.6, 1 mM EDTA, 4 mM MgCl_2_, 2 mM 2-mercaptoethanol). Protein concentration was determined using Nanodrop.

All MBP-fused Yta7-BIM and Yta7-BRD plasmids were transformed into *E. coli* BL21 (DE3) cells, and the transformants were grown in 2 L of LB medium at 37°C. When cell density reached an OD_600_ of 0.8, we induced the expression of the MBP-fused truncated proteins by adding 0.2 mM isopropyl-β-D-thiogalactopyranoside (IPTG) and continued the culture for 12 h at 16°C. We then collected the cells, resuspended in buffer A (20 mM HEPES pH 7.6, 150 mM NaCl, and 10% glycerol), and lysed the cells with a homogenizer (SPX Corporation). The lysate was centrifuged at 20,000x g for 40 min, and the supernatant was collected and loaded into a 5- ml MBPTrap HP column (Cytiva). The MBP fusion protein was eluted using buffer A plus 10 mM maltose and was further purified by size exclusion chromatography through a Superdex 200 column (GE Healthcare) in buffer A.

We transformed the modified pET29a-YS14 plasmids into the Rosetta 2 *E. coli* cells (Novagen) to make the yeast histone proteins. The transformant was inoculated into 6 L of LB medium and grown at 37°C to an OD_600_ of 0.6, then 0.3 mM IPTG was added to induce histone overexpression. The cell culture continued at 37°C for 4 h post induction. Then, we harvested the cells resuspended in 240 ml (40 ml per 1L *E. coli* culture) ice-cold high salt buffer (2 M NaCl, 25 mM HEPES pH 7.5, 10% glycerol, 1 mM 2-mercaptoethanol). Cells were lysed in the homogenizer and the lysate was clarified by centrifugation at 40,000x g for 1 h. We included 50 mM imidazole in the supernatant and loaded the solution into a FPLC with a 5-mL Ni-NTA affinity column. We washed the column with 20 CV of the high salt buffer, then eluted the protein by an imidazole gradient from 20 mM to 500 mM. The eluted peak was collected, concentrated, and run through the HiTrap Heparin HP column (Cytiva) in the high salt buffer to separate the soluble histone octamer. The corresponding fractions were concentrated to 2 mg/ml and the sample quality was examined by SDS-PAGE gel.

### AlphaScreen assay

We performed the AlphaScreen assays using a 6xHis detection kit (PerkinElmer). The biotinylated unmodified and the all-four-lysine acetylated H3 peptides (1-22 aa) were synthesized by GenScript. We individually mixed the purified 50 nM 6xHis-Yta7, 6xHis-MBP- BIM, 6xHis-MBP-BRD, and the complex of 6xHis-MBP-BRD and MBP-BIM with 50 nM biotinylated and unmodified or acetylated H3 (1-22 aa) peptides, respectively, and incubated the mixtures for 1.5 h (for H3 peptide and Yta7 binding) or 2.5 h (for Yta7 domain binding with H3 peptide) on ice with the streptavidin-coated donor beads (5 μg/ml) and Ni-chelated acceptor beads (5 μg/ml) in 50 mM MOPS pH 7.4, 100 mM NaCl, and bovine serum albumin (0.1 mg/ml). We incubated the BRD and BIM mixture for 2.5 h to form complex. The donor beads contain a photosensitizer that converts ambient oxygen into short-lived singlet oxygen upon 680 nm light activation. When the acceptor beads are brought close enough to the donor beads by interaction between 6xHis-tagged proteins and biotinylated peptides, singlet oxygen can diffuse from the donor to the acceptor beads to transfer the energy to the thioxene derivatives of the acceptor beads, resulting in light emission at 520 to 620 nm range.

### Yeast nucleosome core particle reconstitution

We used the 167-bp Widom 601 DNA sequence to reconstitute the yeast nucleosomes. The W601 DNA was amplified by PCR and purified on a HiTrap Q column. After ethanol precipitation, the DNA was resuspended in 25 mM HEPES pH 7.5, 2 M NaCl, 1 mM DTT. The NCPs were reconstituted following the established salt gradient dialysis method ^42^. Briefly, the W601 DNA and purified histone octamer were mixed in a 1.1:1.0 molar ratio, each 100 ul mixture was put in a 0.5-ml Slide-A-Lyzer MINI Dialysis unit (10 kDa cut off, Thermo Scientific), and first dialyzed for 7-8 h into 500 ml buffer A (20 mM Tris-HCl pH 8.0, 1 M NaCl, 1 mM EDTA, 1 mM DTT) at 4°C. The units were then dialyzed into 500 ml buffer B (20 mM Tris-HCl pH 8.0, 0.6M NaCl, 1 mM EDTA, 1 mM DTT) overnight. Finally, the dialysis units were moved into a fresh low salt buffer C (20 mM Tris-HCl pH 8.0, 50 mM NaCl, 1 mM EDTA, 1 mM DTT) and dialyzed for additional 2 - 3 h. The reconstitution results were analyzed on 6% native PAGE and the nucleosome sample was concentrated to 1 mg/ml for future use.

### Sample preparation for cryo-EM

Purified Yta7 at ∼10 mg/mL was incubated with either 2 mM ADP or 2 mM ATPγS plus unmodified H3 peptides (1-24 aa) (GenScript) in 2-fold molar excess to Yta7 hexamer. The mixtures were subsequently incubated at 4°C for 2 h. We added 0.025% octyl β-D-glucoside into the mixture right before cryo-EM grid preparation, which alleviated the preferred particle orientation issue. We applied 3 µL sample droplets two times to freshly glow-discharged (30 s) holey carbon grids (Quantifoil R 2/1 Copper, 300 mesh). EM grids were blotted by filter paper for 3 s after each sample application with the blotting force set to 3. The blotted EM grids were plunge-frozen in liquid ethane cooled by liquid nitrogen in a FEI Vitrobot Mark IV with the sample chamber set to 8°C and 100% humidity.

To maintain the complex assembly, we used the established gradient fixation (GraFix) method to prepare the Yta7-nucleosome complex ^39^. We incubated 2 μM Yta7 with 3 μM nucleosome and 2 mM ATPγS for 1 h on ice in a GraFix buffer (20 mM HEPES, pH 8.0, 150 mM NaCl, 1 mM MgCl_2_, 1 mM DTT), then loaded the sample onto a 12-ml linear 10–30% (v/v) glycerol gradient tube in the GraFix buffer supplemented with 0–0.15% EM-grade glutaraldehyde, and centrifuged the gradient tube at 4°C for 16 h at 35,000 rpm in an SW41 rotor (Beckman Coulter). The sample was fractionated from the bottom to top using an ÄKTA start chromatography system (GE Healthcare Life Sciences). Peak fractions were analyzed by SDS– PAGE, and the fractions containing the desired complexes were pooled and buffer exchanged to remove the glycerol and concentrated to approximately 0.5 mg/ml. For cryo-EM, we applied 3 ul cross-linked Yta7-nucleosome complex onto freshly glow-discharged (30 s) holey carbon grids (Quantifoil R 2/1 Gold, 300 mesh) and blotted for 3 s with a piece of filter paper. The grids were vitrified by plunging them into liquid ethane in a FEI Vitrobot Mark IV set to 8°C and 100% humidity. The cryo-EM grids were stored in liquid nitrogen until data collection.

### Cryo-EM data collection

Cryo-EM datasets were collected on a TFS Titan Krios electron microscope operated at 300 kV and equipped with a K3 summit direct electron detector (Gatan). All EM images were recorded with the objective lens defocus range of –1.0 to –2.0 μm at a nominal magnification of 105,000× using SerialEM ^43^, with an effective calibrated image pixel size of 0.414 Å. The camera was operated in the super-resolution counting and movie mode with a dose rate of 0.88 electrons per Å^2^ per second per frame. A total of 75 frames were recorded for each micrograph stack.

### Image processing

The dataset of ADP-bound Yta7 contained 12,456 movie stacks. The full dataset was split into five subgroups and imported to Relion-3.1 ^44^. Micrographs in each subgroup were drift-corrected with electron-dose weighting and 2x binned using MotionCor2 (version 1.3.2) ^45^, resulting in a pixel size of 0.828 Å at sample level. The defocus value of each micrograph was estimated by CTFFIND4 ^46^. We first manually picked a small particle dataset to generate templates that were subsequently used for autopicking in Relion 3.1 ^44^. A total of 3,130,724 particles were picked and extracted with 4x binning. These particles were imported into Cryosparc2 (version 3.2.0) to perform several rounds of 2D classifications and create a starting 3D model ^47^. The selected good 2D classes contained 593,691 particle images. These particle images were converted to the RELION format using UCSF PyEM (version 0.5) (https://github.com/asarnow/pyem), and further processed with 3D classification and refinement in Relion-3.1. Two major 3D classes with 299,123 and 215,755 particles, respectively, were combined to generate a 3D map at 3.3 Å resolution. After further per-particle CTF refinement and Bayesian polishing, the final 3D map reached an overall resolution of 3.1 Å, based on the gold-standard Fourier shell correlation (FSC) curve of two half maps at the standard threshold of 0.143 ^48^.

The particle images that were used for final 3D reconstruction of the 3D map of the ADP-bound Yta7 complex were further processed using CryoDRGN ^50^. The structural heterogeneity of the dataset was analyzed, and the particles were separated into 20 clusters using k-mean clustering method. Structural analysis showed that the clusters differed from each other by the number of BRD/BIM domains that can be resolved on top of the AAA+ domains, with three BRD/BIM domains being the most resolved in the spiral. The corresponding particles producing three stable BRD/BIM domains were then used to run another round of refinement in cryoSPARC, leading to the final resolution of 3.9 Å.

The dataset of the ATPγS and H3 tail-bound Yta7 contained 12,085 movie stacks. These images were processed similarly as described above. A total of 4,062,171 particles were picked. The selected good 2D classes contained 572,451 particle images. After 3D classification Relion- 3.1, one 3D class with 431,065 particle images was selected for further refinement, leading to a 3D map with bound H3 peptide at an estimated resolution of 3.3 Å. Further per-particle CTF refinement and Bayesian polishing improved the map to 3.0 Å. A second 3D class with 109,554 particle images was also selected for refinement, leading to a 3D map at 5.6 Å resolution with a spirally arranged top BRD-tier. Local resolution 3D maps were calculated using ResMap ^49^.

The dataset of the Yta7–-nucleosome complex contained 14,937 movie stacks and was processed mostly by using Cryosparc2 as described above (**Supplementary Fig. 9**). Autopicked particle dataset with 3,152,817 particle images went through several rounds of 2D class averages in which the Yta7 region was well defined, but the top nucleosome region was blurry. We selected a dataset of 895,586 particle images with 2D classes that had relatively solid nucleosome density. After ab initio 3D reconstruction and 3D classification, we identified one 3D class map with some nucleosome density; this class had a total of 213,712 particle images. We downscaled the particle images by a factor of 4 to the pixel size of 3.312 Å, and refined a 3D map; however, the nucleosome was still not well resolved in the map. Further heterogenous refinement also did not improve the nucleosome density. We next performed three-dimensional variability analysis (3DVA) implemented in cryoSPARC to visualize the dynamics of the nucleosome in complex with Yta7. The results showed that nucleosome binding was highly variable. Among the 3D frames of the 3DVA analysis, we identified one 3D map in which the nucleosome position is resolved. This map had an estimated resolution of 18 Å based on the gold-standard Fourier shell correlation (FSC) 0.143 criterion ^48^. Finally, we analyzed the Yta7 conformation in the presence of nucleosome, by performing the focused refinement of downscaled particle images of Yta7-nucleosome complex by excluding the nucleosome region. This led to a 14-Å resolution 3D map of Yta7 in which the BRD/BIM-tier formed a flat ring above the AAA1-tier.

### Atomic model building and refinement

We first used the Robetta server ^51^ to generate a homology model for Yta7 based on the pombe Abo1 structure (PDB 6JQ0). The starting Yta7 model was split into the AAA1, AAA2, and the C-terminal helical regions, and docked individually into the cryo-EM maps using UCSF Chimera ^52^. We then manually adjusted and built missing residues in our high-resolution cryo-EM map in Coot ^53^. The resulting model was subjected to several iterations of real-space refinement in PHENIX ^54^ and manual adjustment in Coot ^53^. Both the original maps and the maps sharpened by deepEMhancer were used to refine most of the atomic models ^55^. We used the UCSF ChimeraX to make the figures ^56^. For the low-resolution 3D map of the Yta7-nucleosome, we performed rigid-body docking with the yeast nucleosome structure (PDB 6GEJ) and our atomic Yat7 model. The statistics of model refinement are shown in **Supplementary Table 2**.

---

**Supplementary Figure 1.**
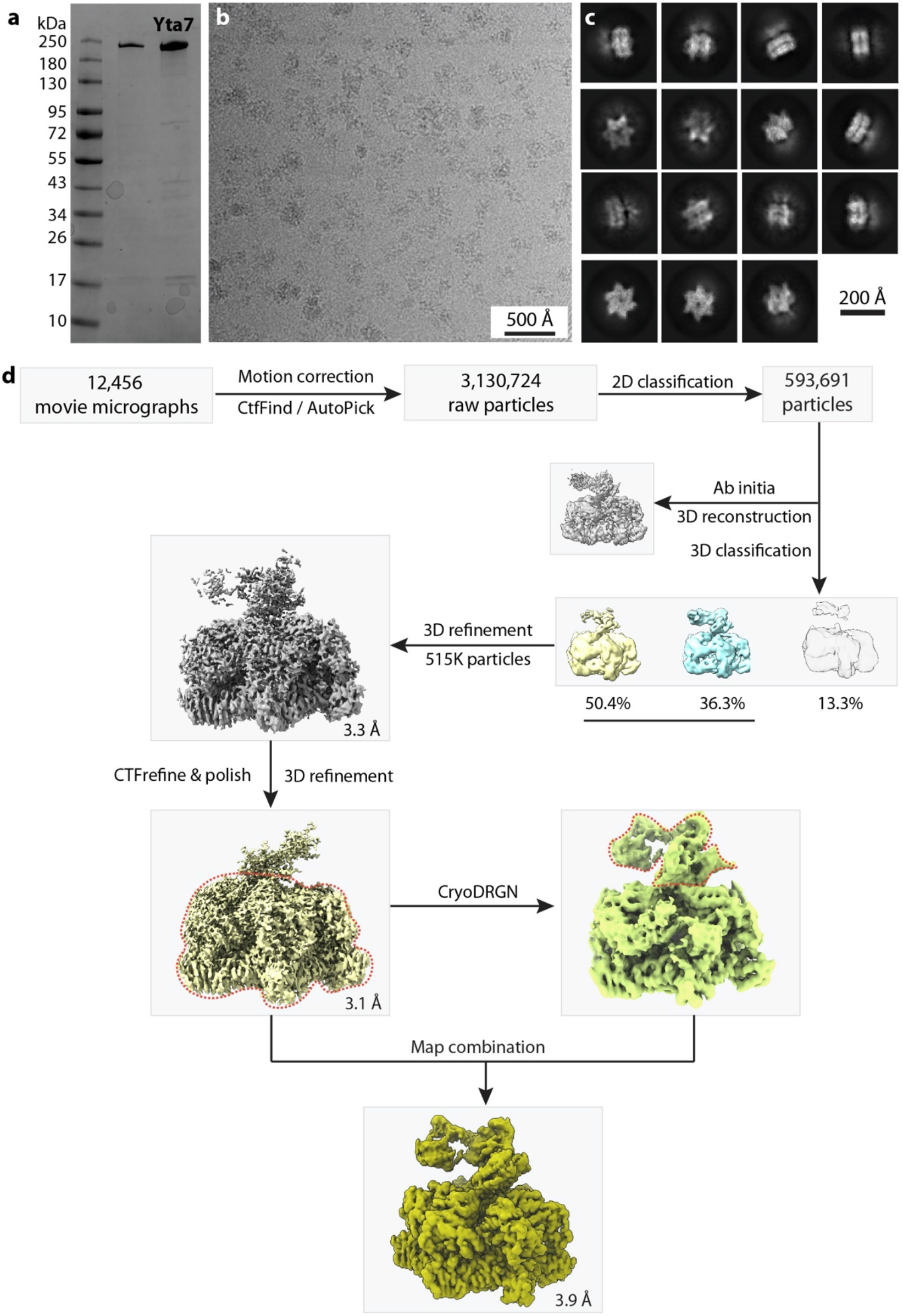
Workflow of cryo-EM data processing and 3D reconstruction of Yta7 in the ADP state. **a**, SDS-PAGE gel of purified Yta7. **b**, A typical raw micrograph. A total of 12,456 such micrographs were recorded in this study. **c**, Selected 2D class averages. **d**, Workflow of cryo-EM data processing of purified Yta7 in the presence of ADP.

**Supplementary Figure 2.**
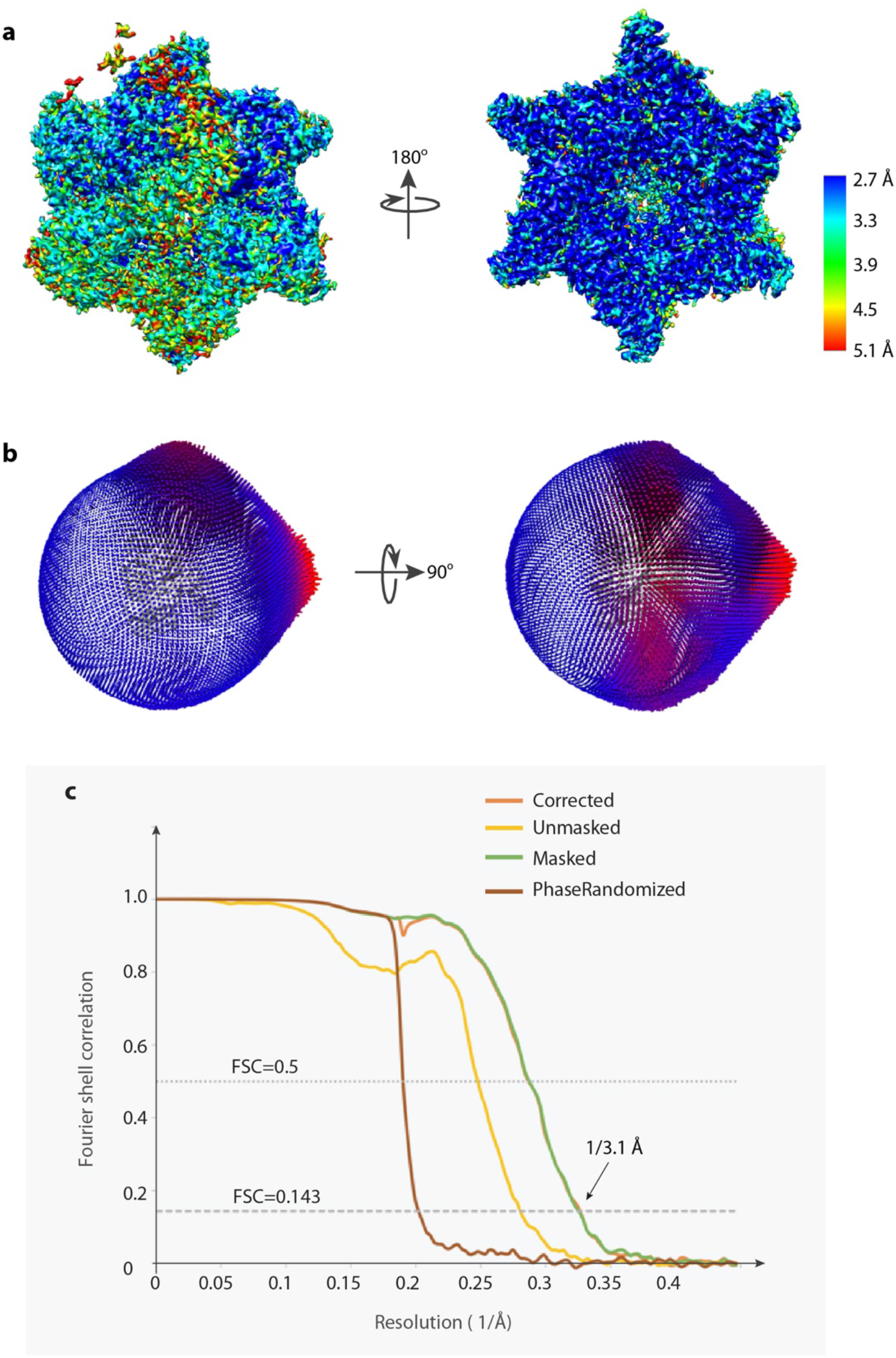
Resolution estimation of the 3D map of the ADP-bound Yta7 hexamer. **a**, Color coded local resolution map of the ADP-bound Yta7 EM map. **b**, Angular distribution plot of particles used in the final 3D reconstruction. **c**, The Fourier shell correlation curves of the final 3D map.

**Supplementary Figure 3.**
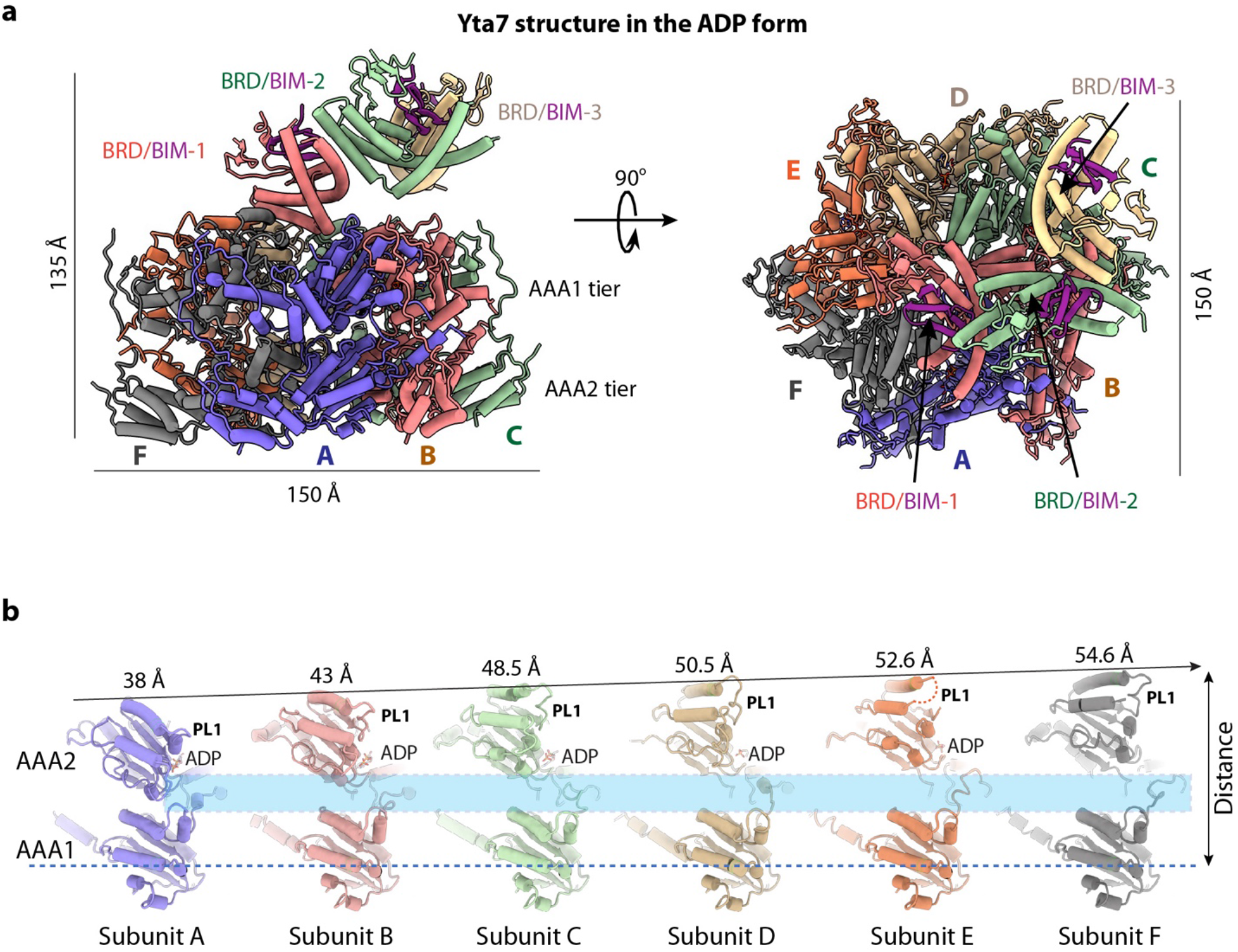
Structure of the ADP-bound Yta7 hexamer. **a**, A side and a top view of the ADP-bound Yta7 structure in cartoon. Subunits are individually colored. The three resolved BRD domains are in orange, dark green, and salmon, respectively, and their associated BIM motifs in magenta. Dimensions of the structure are shown in Å. **b**, The six Yta7 subunits are aligned with their respective AAA2 domains and shown individually in the same orientation. The distance between α2 helix of AAA1 and α3 helix of AAA2 in each subunit is labeled on top, revealing a spiral arrangement of the AAA1 domains. The cyan rectangle marks the variably configured linkers between AAA1 and AAA2.

**Supplementary Figure 4.**
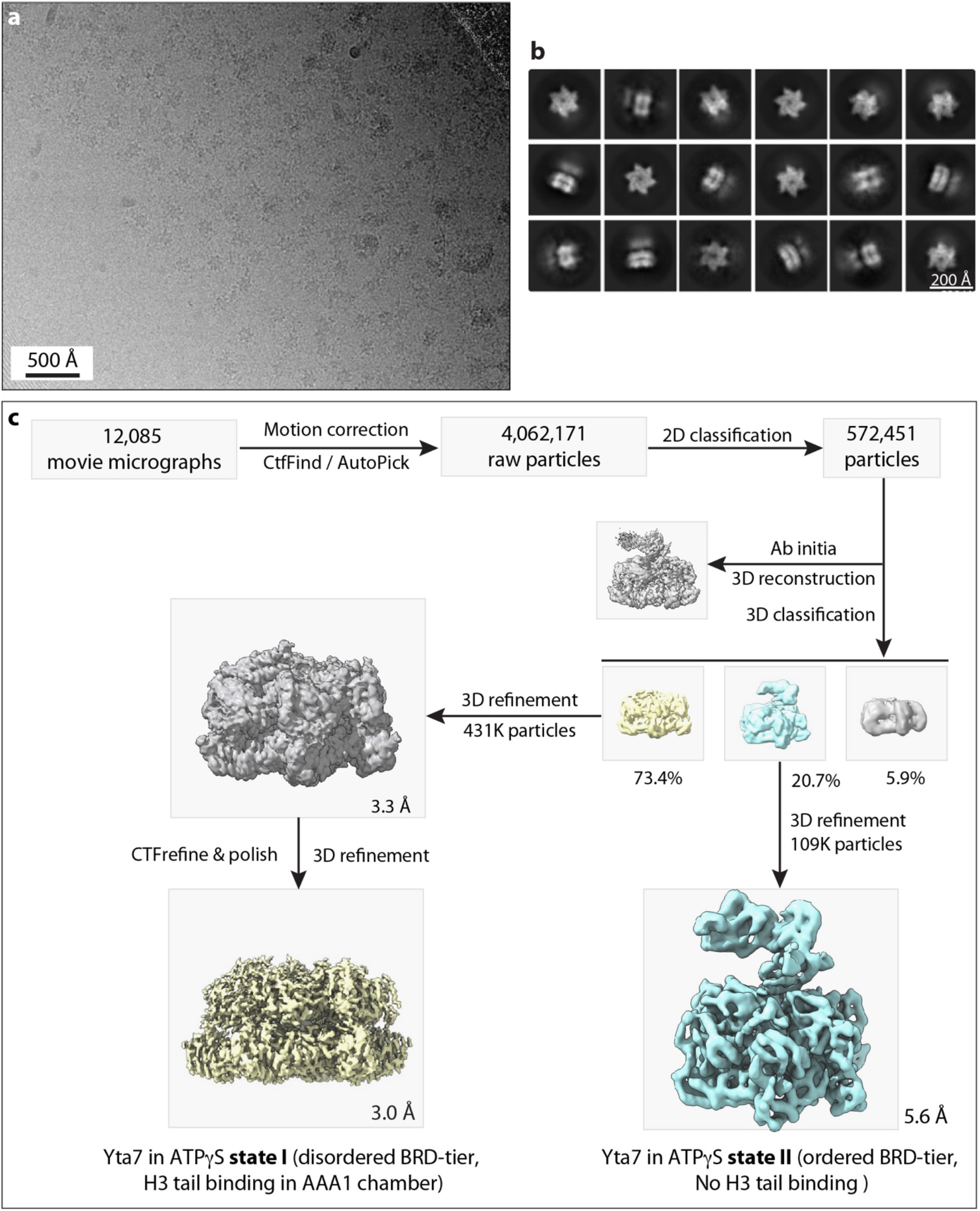
Workflow of cryo-EM data processing and 3D reconstruction of Yta7 bound to ATPγS and an H3 peptide. **a**, A typical raw micrograph. A total of 12,085 such micrographs were recorded in this study. **b**, Selected 2D class averages. **c**, Workflow of cryo-EM data processing using Relion and cryoSPARC.

**Supplementary Figure 5.**
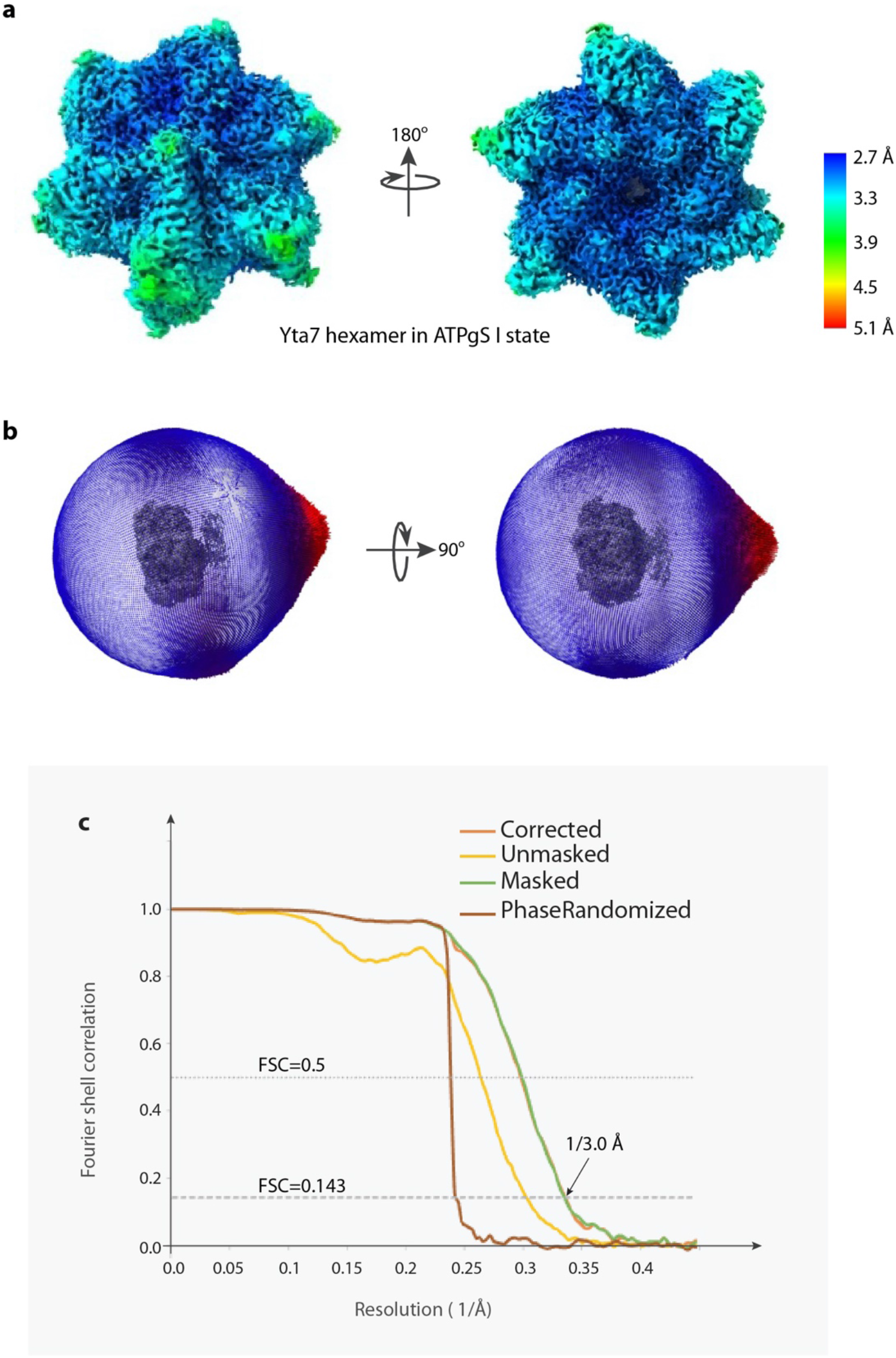
Resolution estimation of the 3D reconstruction of Yta7 bound to ATPγS and an H3 peptide. **a**, Colored coded local resolution map of the 3D reconstruction of the ATPγS-bound Yta7 in state I (bound to H3 tail). **b**, Angular distribution plot of particles used for the final 3D refinement and reconstruction. **c**, Fourier shell correlation curves of the final 3D map suggests an average resolution of 3.0 Å.

**Supplementary Figure 6.**
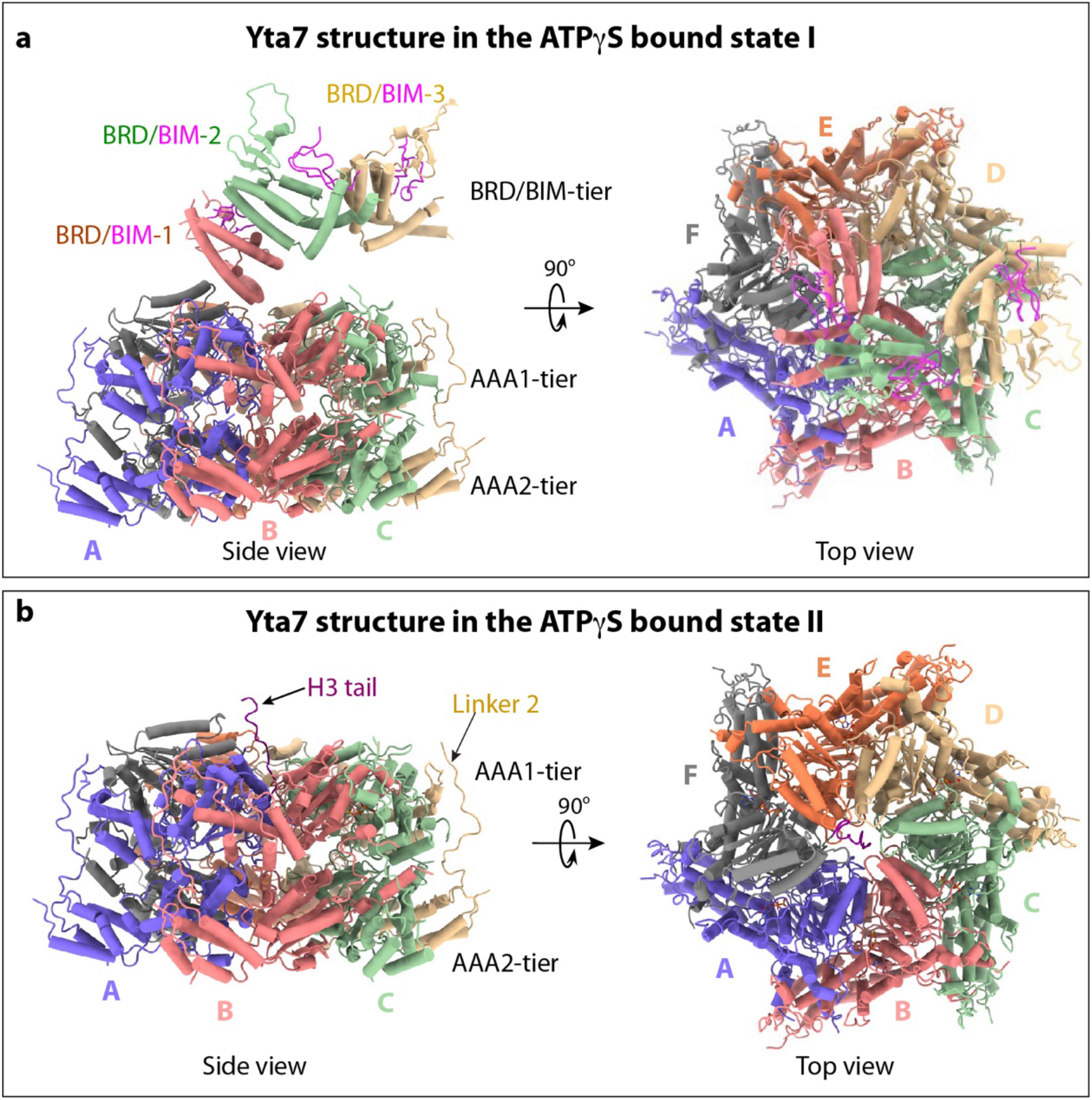
Structural comparison of the ATPγS-bound Yta7 hexamer in states I and II. **a**, Cartoon views of the structure of ATPγS-bound Yta7 in state I in which the top BRD/BIM-tier is partially ordered, with the three BRD/BIM resolved and modeled. No H3 peptide was observed in this state, because the proximal BRD/BIM-1 blocks the peptide entry to the central chamber. **b**, Cartoon views of the Yta7 structure bound to ATPγS and H3 peptide (state II). Subunits are individually colored. The BRD/BIM-tier on top is disordered in this state such that the Yta7 substrate entry is unblocked and H3 tail is partially resolved.

**Supplementary Figure 7.**
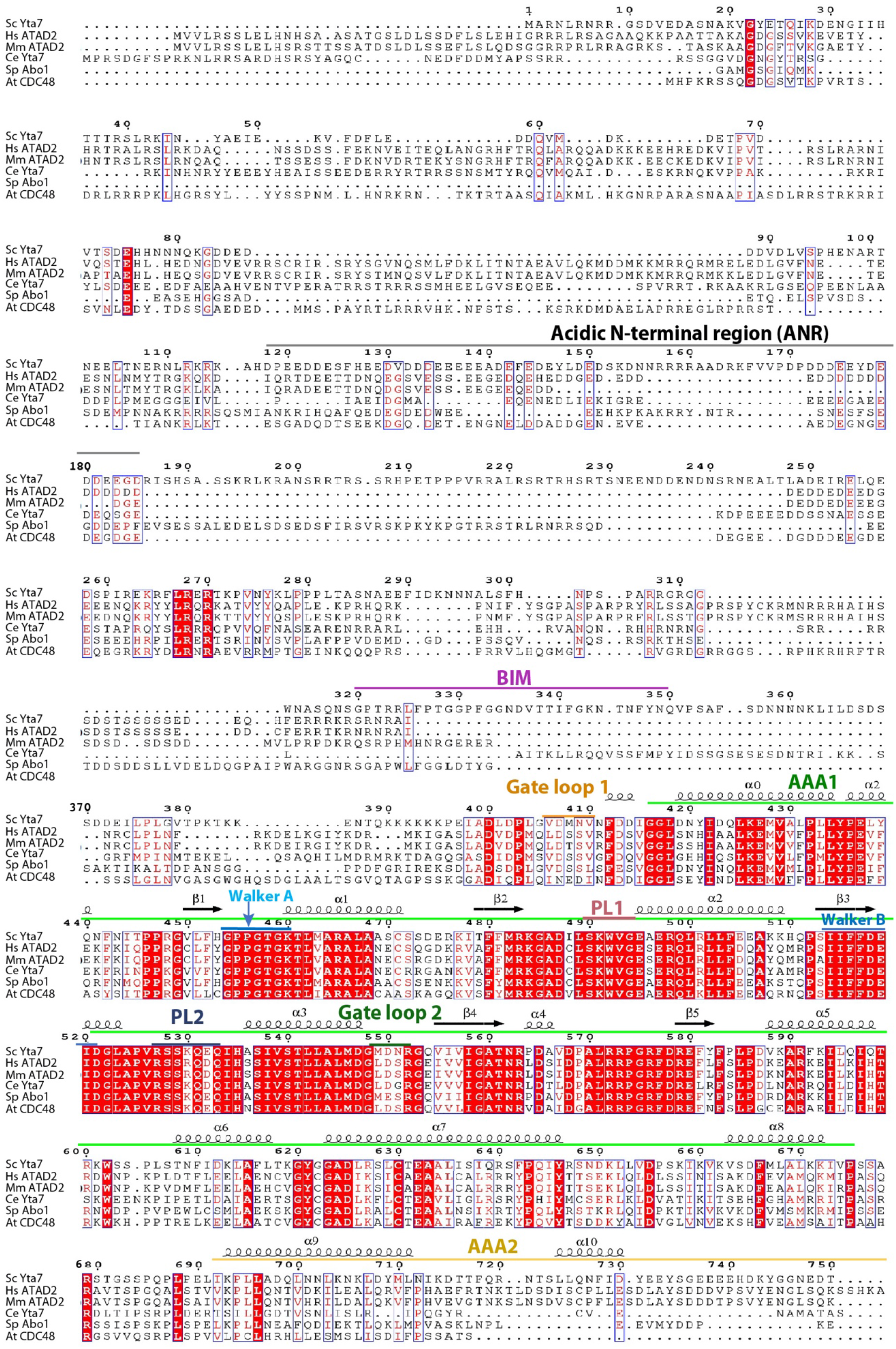
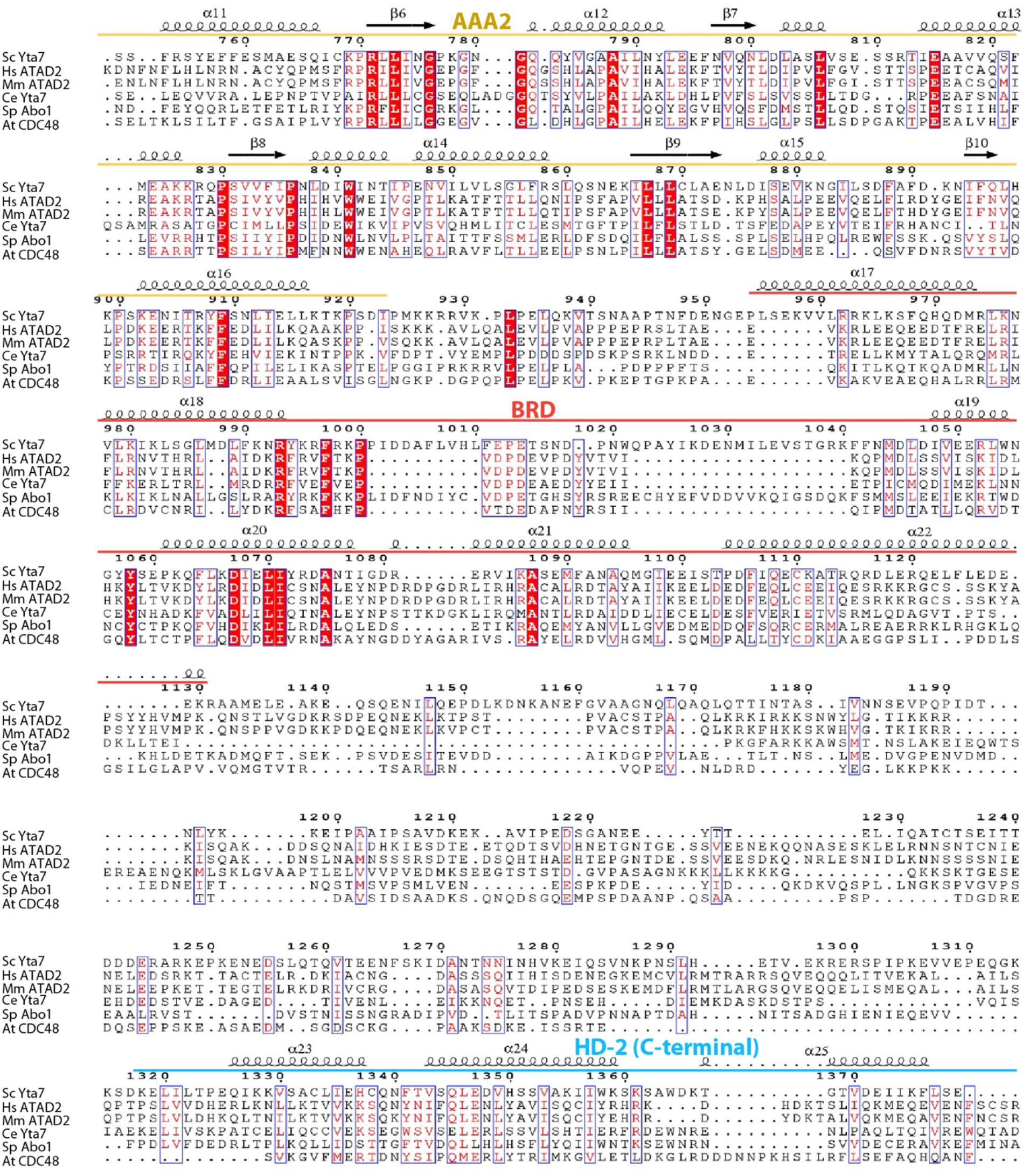
Sequence alignment of six Yta7 homologues. The amino acid sequences of *S. cerevisiae* Yta7, *H. sapiens* ATAD2, *M. musculus* ATAD2, *C. elegans* Yta7, *S. pombe* Abo1, and *A. thaliana* CDC48 were aligned in ClustalW and further analyzed by ESPript3.0. Major domains and key regions in Sc Yta7 are labeled above the sequence.

**Supplementary Figure 8.**
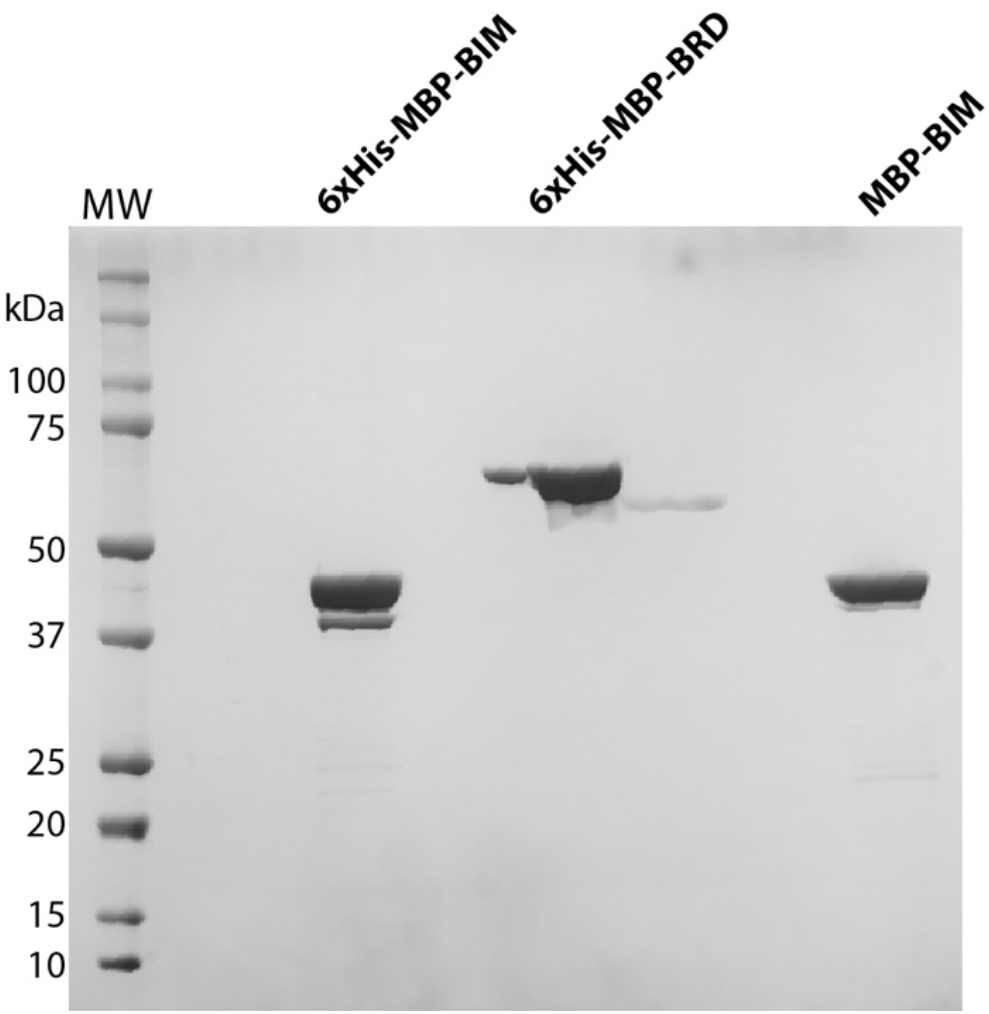
SDS-PAGE gel of purified 6xHis-MBP-BIM, 6xHis-MBP-BRD, and MBP-BIM. These truncated Yta7 domain proteins were used in the AlphaScreen assay. MW: molecular weight marker.

**Supplementary Figure 9.**
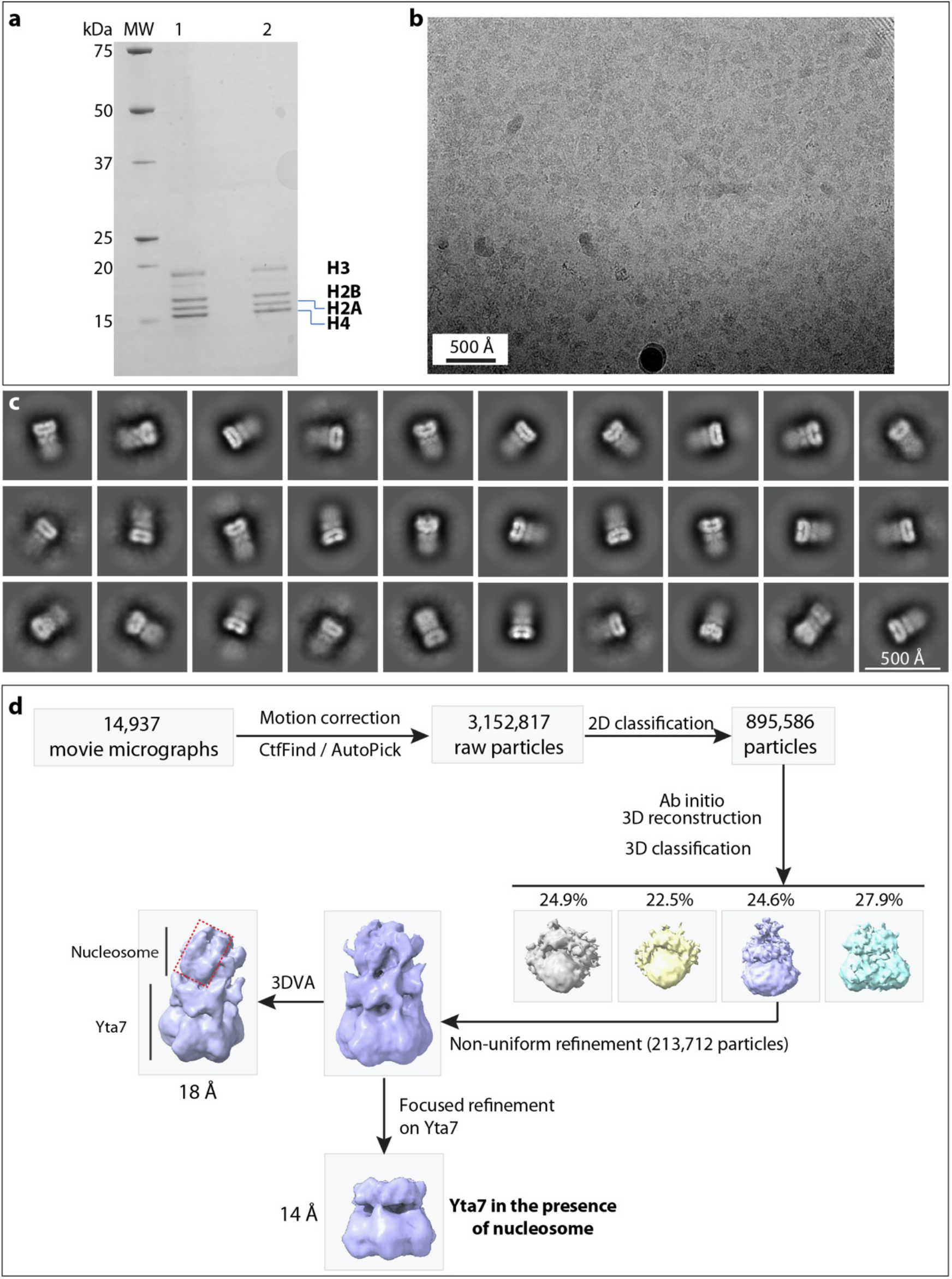
Cryo-EM analysis of the Yta7-nucleosome complex. **a**, SDS-PAGE gel of purified yeast histone octamer. **b**, A typical raw micrograph of the Yta7– nucleosome complex bound to ATPγS. A total of 14,937 micrographs were recorded. **c**, Selected 2D class averages. The fuzzy density associated with the well-ordered Yta7 density is from the bound nucleosome. **d**, Workflow of data processing in cryoSPARC.

**Supplementary Table 1.**
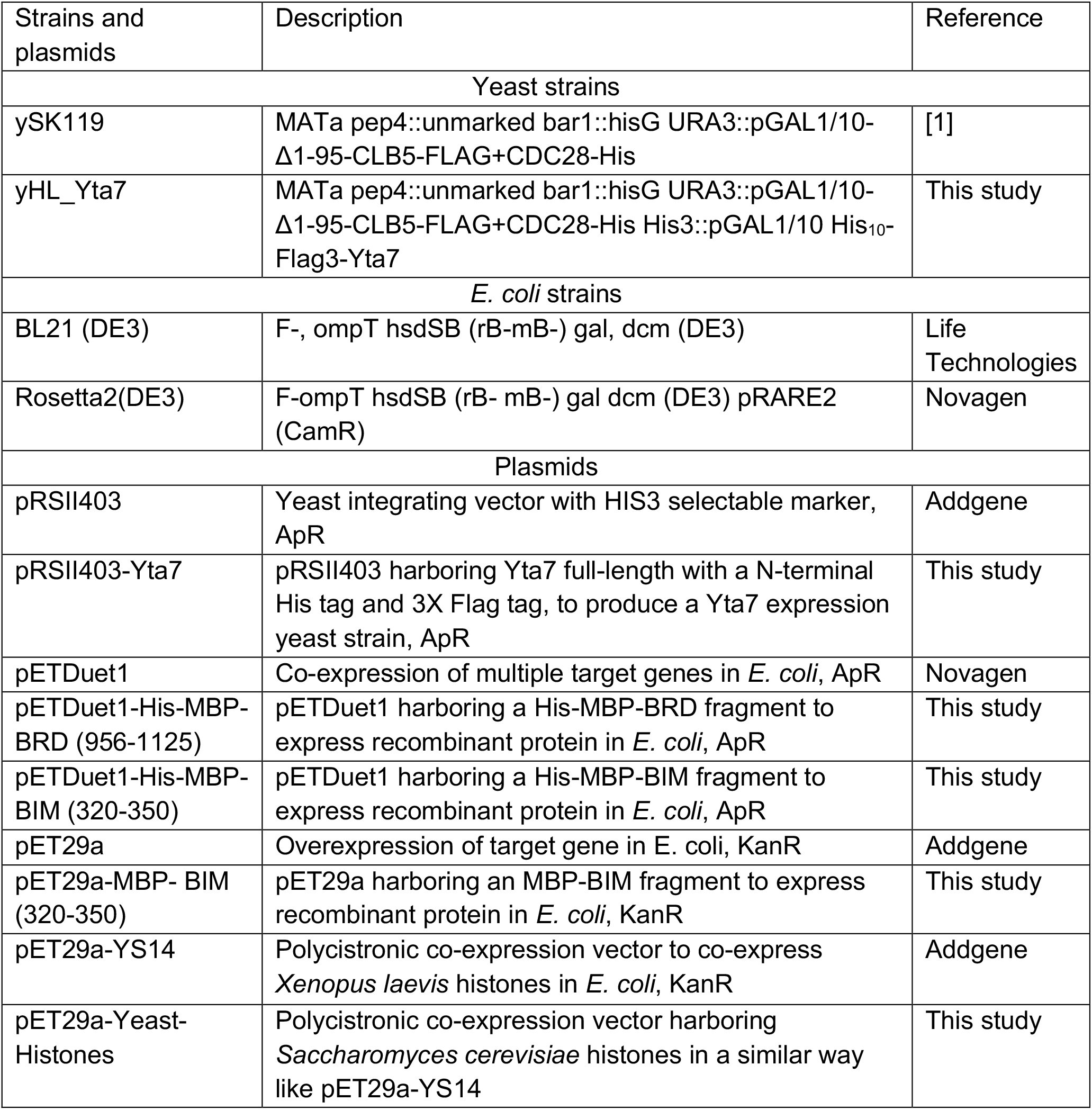
List of strains and plasmids used in this study.

**Supplementary Table 2.**
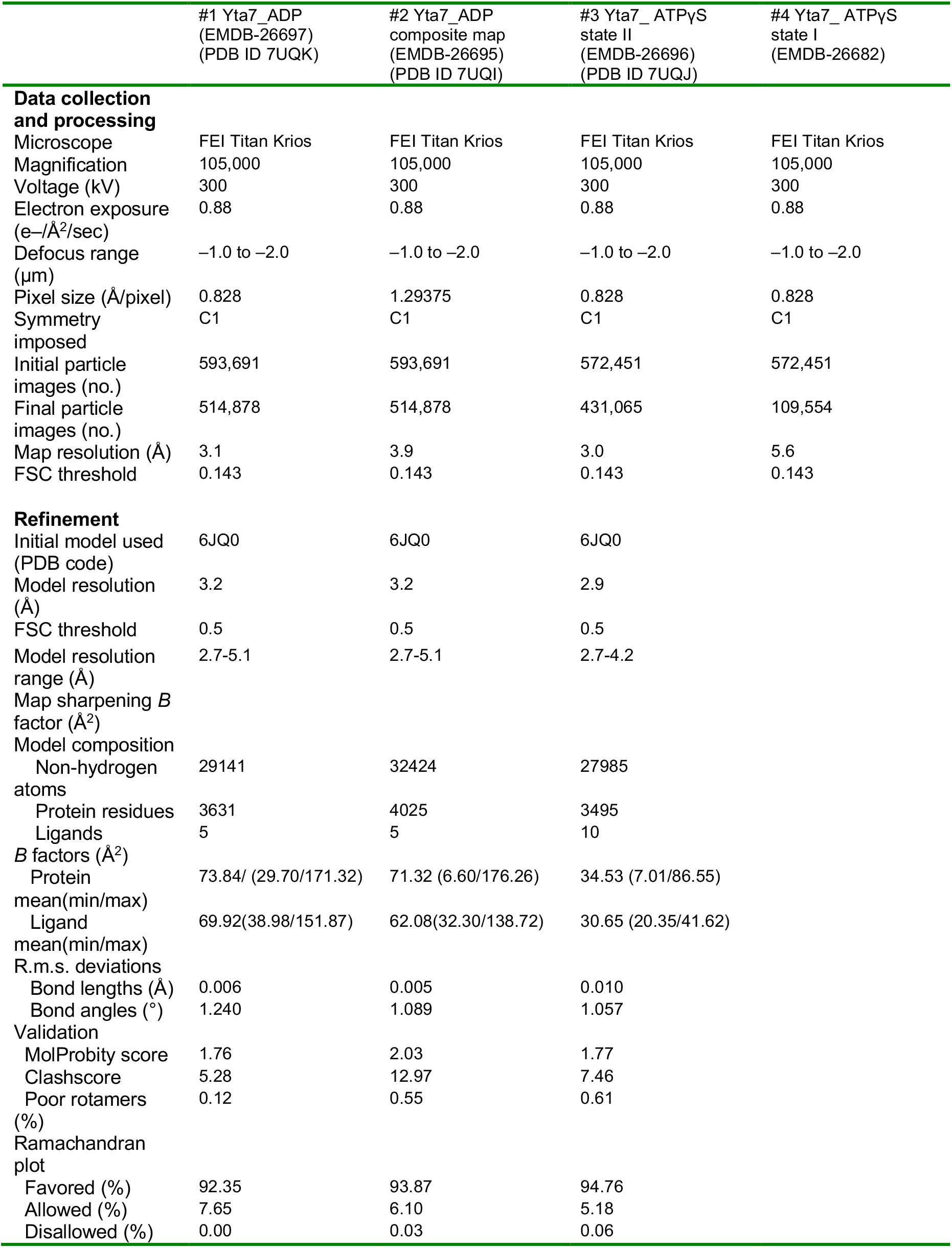
Cryo-EM data collection, refinement, and validation statistics.

## Notes

### Competing Interest Statement

The authors have declared no competing interest.

